# Integration of metabolomic and transcriptomic analyses reveals novel regulatory functions of the ChREBP transcription factor in energy metabolism

**DOI:** 10.1101/2024.09.17.613577

**Authors:** Jie An, Inna Astapova, Guofang Zhang, Andrew L. Cangelosi, Olga Ilkayeva, Hannah Marchuk, Michael J. Muehlbauer, Tabitha George, Joseph Brozinick, Mark A. Herman, Christopher B. Newgard

## Abstract

Carbohydrate Response Element-Binding Protein (ChREBP) is a transcription factor that activates key genes involved in glucose, fructose, and lipid metabolism in response to carbohydrate feeding, but its other potential roles in metabolic homeostasis have not been as well studied. We used liver-selective GalNAc-siRNA technology to suppress expression of ChREBP in rats fed a high fat/high sucrose diet and characterized hepatic and systemic responses by integrating transcriptomic and metabolomic analyses. GalNAc-siChREBP-treated rats had lower levels of multiple short-chain acyl CoA metabolites compared to rats treated with GalNAc-siCtrl containing a non-targeting siRNA sequence. These changes were related to a sharp decrease in free CoA levels in GalNAc-siChREBP treated-rats, accompanied by lower expression of transcripts encoding enzymes and transporters involved in CoA biosynthesis. These activities of ChREBP likely contribute to its complex effects on hepatic lipid and energy metabolism. While core enzymes of fatty acid (FA) oxidation are induced by ChREBP knockdown, accumulation of liver acylcarnitines and circulating ketones indicate diversion of acetyl CoA to ketone production rather than complete oxidation in the TCA cycle. Despite strong suppression of pyruvate kinase and activation of pyruvate dehydrogenase, pyruvate levels were maintained, likely via increased expression of pyruvate transporters, and decreased expression of lactate dehydrogenase and alanine transaminase. GalNAc-siChREBP treatment increased hepatic citrate and isocitrate levels while decreasing levels of distal TCA cycle intermediates. The drop in free CoA levels, needed for the 2-ketoglutarate dehydrogenase reaction, as well as a decrease in transcripts encoding the anaplerotic enzymes pyruvate carboxylase, glutamate dehydrogenase, and aspartate transaminase likely contributed to these effects. GalNAc-siChREBP treatment caused striking increases in PRPP and ZMP/AICAR levels, and decreases in GMP, IMP, AMP, NaNM, NAD(P), and NAD(P)H levels, accompanied by reduced expression of enzymes that catalyze late steps in purine and NAD synthesis. ChREBP suppression also increased expression of a set of plasma membrane amino acid transporters, possibly as an attempt to replenish TCA cycle intermediates. In sum, combining transcriptomic and metabolomic analyses has revealed regulatory functions of ChREBP that go well beyond its canonical roles in control of carbohydrate and lipid metabolism to now include mitochondrial metabolism and cellular energy balance.

## Introduction

Excess consumption of diets enriched in fats and sugars leads to obesity and a myriad of metabolic derangements, including insulin resistance, glucose intolerance, type 2 diabetes, and elevated tissue and plasma lipid levels. The effects of consumption of high-calorie diets on intermediary metabolism are orchestrated in part by transcription factors, including carbohydrate response element binding protein (ChREBP), sterol regulatory element-binding protein (SREBP), and members of the peroxisome proliferator-activated receptor (PPAR) family and its coactivators such as PGC-1, each with the capacity to regulate genomic programs affecting diverse metabolic functions.

Our group has focused on the metabolic regulatory roles of ChREBP in the liver during feeding of high calorie diets^1–5^. Our prior studies and those of others have demonstrated that ChREBP is strongly activated by diets enriched in sucrose or fructose, resulting in induction of genes involved in glucose and fructose catabolism such as the GLUT2 glucose transporter and the GLUT5 fructose transporter, ketohexokinase (KhK), glucokinase regulatory protein (Gckr), triose kinase (Tkfc), glucose-6- phosphatase (G6pc), and pyruvate kinase liver/red blood cell isoform (PKLR) as well as genes involved in synthesis of lipids from glucose (*de novo* lipogenesis, DNL) such as ATP-citrate lyase (ACLY) and fatty acid synthase (FASN)^1–11^. Our recent work demonstrates that ChREBP also regulates genes involved in metabolism of branched-chain amino acids (BCAA) in liver^12^. Re-feeding of fasted rats with a diet high in fructose caused a large increase in expression of ChREBP, paralleled by increased expression of the transcript encoding branched-chain ketoacid dehydrogenase kinase (BCKDK), the enzyme that phosphorylates and inhibits activity of branched-chain ketoacid dehydrogenase (BCKDH), and a decrease in expression of PPM1K, the phosphatase that dephosphorylates and activates BCKDH. Further, adenovirus-mediated overexpression of ChREBP in rat liver was sufficient to cause an increase in BCKDK and decrease in PPM1K transcript levels^12^.

These recent findings suggest that ChREBP may regulate cellular and systemic metabolism in ways that transcend its well-studied effects on glucose and lipid metabolism to include hepatic amino acid metabolism and other pathways. Also suggestive of alternate functions is the finding that despite potent activation of glycolytic flux by ChREBP, the liver is reported to shift from fatty acid oxidation to pyruvate and lactate oxidation in response to global ChREBP knockout^13^. This was associated with decreases in cytosolic [NAD+]/[NADH] and [ATP]/[ADP] [Pi] ratios and suppression of flux through the anaplerotic enzyme pyruvate carboxylase^13^. However, the mechanisms and specific ChREBP transcriptional targets that might lead to these changes were not identified. Further studies to understand broader metabolic regulatory effects of ChREBP therefore seem warranted.

In the present study we used ChREBP-specific siRNAs conjugated with GalNAc to suppress expression of ChREBP in liver of rats fed a high fat/high sucrose (HF/HS) diet, and then performed extensive metabolomic and transcriptomic profiling to gain insight into currently unknown mechanisms by which ChREBP regulates cellular metabolism. The GalNAc conjugation method was chosen for its liver-selective transgene delivery and long-term modulation of gene expression^14^. Using this technology, we achieved robust suppression of ChREBP expression in liver for 4 weeks, allowing us to integrate comprehensive metabolomic and transcriptomic profiles to provide a broad overview of the role of ChREBP in integration of hepatic fuel metabolism. This approach revealed novel effects of ChREBP knockdown on hepatic metabolic pathways, including: 1) a sharp decrease in free CoA levels, accompanied by reduced expression of genes involved in CoA biosynthesis and decreases in levels of multiple short-chain acyl CoA metabolites; 2) increased expression of core fatty acid oxidation enzymes accompanied by increased hepatic acylcarnitine and circulating ketone levels, suggestive of incomplete fatty acid oxidation; 3) coordinated regulation of pyruvate metabolism, including upregulation of plasma membrane pyruvate transporters, decreased expression of LDHa and alanine transaminase, and decreases in expression of pyruvate carboxylase and PDK-4 to favor PDH-catalyzed oxidative metabolism of pyruvate; 4) upregulation of multiple plasma membrane amino acid transporters, possibly representing an attempt to compensate for decreased anaplerosis from carbohydrates; and 5) reduced expression of ATIC and HPRT, encoding the last steps in the *de novo* and salvage pathways of purine synthesis, respectively, and QPRT, which catalyzes a late step in the *de novo* pathway of NAD synthesis, accompanied by decreases in GMP, IMP, AMP, NaMN, NAD(P), and NAD(P)H levels. In aggregate, our findings define new roles for ChREBP in integration of diverse pathways of cellular fuel metabolism and energy balance extending well beyond its canonical roles in regulation of glycolytic and lipogenic pathways.

## Results

### GalNAc-siRNA achieves robust knockdown of ChREBP to affect markers of glucose and lipid metabolism

To induce obesity, a cohort of obesity-prone (OP-CD) male Sprague Dawley rats were fed with a high-fat/high sucrose (HF/HS) diet for a period of 10 weeks beginning at 4 weeks of age. After this period of HF/HS feeding, the rats were divided into groups of 5-6 rats, and injected subcutaneously with 1) an siRNA specific for ChREBP conjugated with GalNAc (GalNAc- siChREBP); 2) a control, non-targeting siRNA conjugated with GalNAc (GalNAc-siCtrl); or 3) saline (Sal). Additional doses of each GalNAc-siRNA construct or Sal were injected at 10, 18 and 25 days after the first treatment. An intraperitoneal glucose tolerance test (IPGTT) was performed 20 days after the first injection of the GalNAc constructs or saline. At day 26, rats in all groups received a priming bolus injection of D2O (10 ml/kg), followed by provision of drinking water supplemented with D2O to a level of 4% to allow measurement of hepatic *de novo* lipogenesis (DNL). A blood sample was obtained at day 27 to measure D2O enrichment in plasma water, and all animals were anesthetized and sacrificed 28 days after first injection of the GalNAc constructs or saline for collection of blood and tissues for transcriptomic, immunoblot, DNL, and metabolomic analyses.

Consumption of foods and beverages high in added sugars, and particularly fructose, or conditions like obesity which increase hepatic ‘reductive stress’ and the abundance of hepatic carbohydrate metabolites, activate expression of ChREBP-α, which then acts upon an alternative promoter in the ChREBP gene to activate transcription of ChREBP-β, the more potent transcriptional activator among the two ChREBP isoforms^1,3,9,15^. Injection of GalNAc-siChREBP caused almost complete suppression of ChREBP-α and ChREBP-β transcripts in liver relative to the saline or GalNAc-siCtrl groups (**Figure 1A**) 28 days post-treatment, accompanied by a similarly potent suppression of ChREBP-α protein expression as measured by immunoblot analysis. (**Figure 1B)**. The reduction in ChREBP expression was accompanied by marked reductions in the hepatic expression of canonical ChREBP targets involved in glucose and fructose transport, glycolysis, glucose production, fructolysis, *de novo* lipogenesis, and packaging and export of VLDL lipoprotein particles, consistent with prior studies (**Figure 1C**)^1–11^. Body weights were not different among rats in the 3 main experimental groups over the time course of treatment with GalNAc-siChREBP, GalNAc-siCtrl, or saline (**Supplemental Figure 1A**). An intraperitoneal glucose tolerance test performed at day 20 after onset of the GalNAc-siRNA treatments revealed no significant differences in glucose excursion among the 3 experimental groups (**Supplemental Figure 1B**). Blood samples were also collected at the time of sacrifice (8 days after the IPGTT) and used for measurements of glucose and insulin levels. At that time point, no differences in blood glucose were observed (**Supplemental Figure 1C**), whereas insulin levels were lowered by GalNAc-siChREBP treatment compared to the GalNAc-siCtrl-treated group (**Supplemental Figure 1D**), resulting in a statistically significant decrease in HOMA-IR, a measure of insulin sensitivity, in the GalNAc-siChREBP-treated group (**Supplemental Figure 1E**).

**Figure 1.**
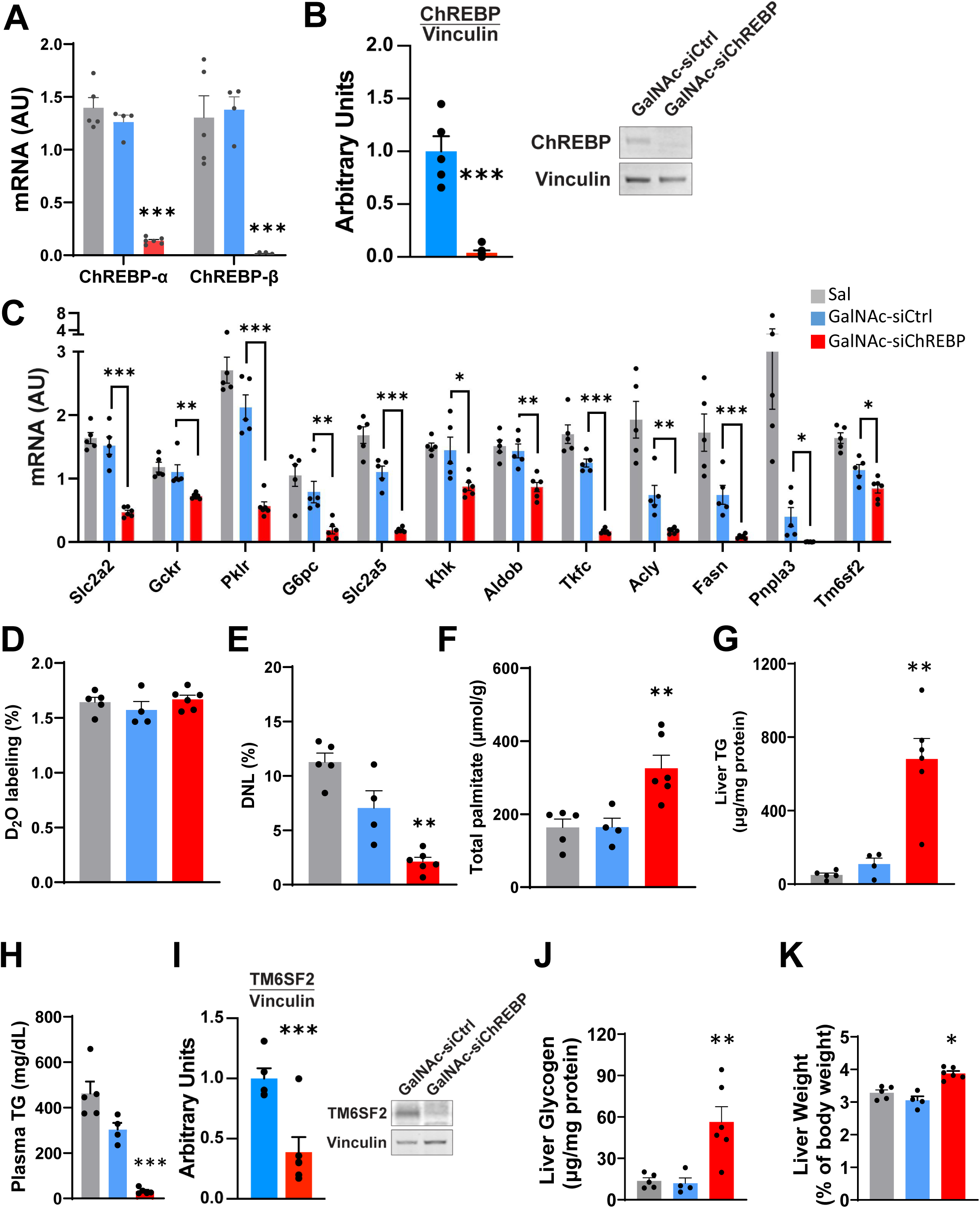
Effects of GalNAc-siRNA treatment on transcript levels. Rats fed on HF/HS diet received multiple injections of GalNAc-siChREBP, GalNAc-siCtrl, or saline over a 28 day period prior to sacrifice and collection of liver samples q-PCR and biochemical analyses **A.** Levels of transcripts encoding ChREBP-alpha and ChREBP-beta isoforms measured by q-PCR. **B.** Levels of ChREBP protein shown by representative immunoblot (left) and densitometric analysis of 5-6 samples per group (right). **C.** Levels of transcripts encoded by known ChREBP target genes including GLUT-2 (Slc2a2), glucokinase regulatory protein (Gckr), pyruvate kinase, liver isoform (Pklr), glucose-6-phosphatase (G6pc), GLUT-5 (Slc2a5), ketohexokinase (Khk), aldolase b (Aldo b), triose kinase (Tkfc), ATP-citrate lyase (Acly), fatty acid synthase (Fasn), patatin-like phospholipase domain-containing protein 3 (Pnpla3), and transmembrane 6 superfamily 2 (Tm6sf2). **D.** All rats received a bolus of deuterium oxide (D2O, 10 ml/kg body weight) 26 days after the initial GalNAc-siRNA or saline injection and then received drinking water supplemented with 4% D2O until day 28, with a blood sample taken at day 27 to measure D2O labeling of plasma water. Panel D demonstrates that D2O enrichment in plasma across the three experimental groups is equivalent. **E.** Percent labeling of newly synthesized palmitate with D2O. **F.** Total liver palmitate content (free and esterified). **G.** Liver triglyceride (TG) content. **H.** Plasma triglyceride levels. **I.** Levels of TM6sf2 protein shown by representative immunoblot (right) and densitometric analysis of 4 samples per group (left). **J.** Liver glycogen levels. **K.** Liver weights. Data are the mean ± standard error of the mean (SEM) for 4-6 independent rats/liver samples for the GalNAc-siChREBP-treated group and 4-5 rats/liver samples for the GalNAc-siCtrl and saline-treated groups. Statistical analysis performed using one-way ANOVA followed by Tukey test (A, C-H, J, and K) or unpaired t-test (B and I). Symbols *, **, and *** indicate that a value was significantly lower in the GalNAc-siChREBP treatment group compared to either the GalNAc-siCtrl or saline groups, with p < 0.05, p < 0.01, and p < 0.005, respectively.

### Effect of ChREBP suppression on DNL, hepatic fuel storage, lipid metabolism, and liver weight

All animal groups received a priming injection of D2O and had D2O added to their drinking water beginning two days before sacrifice to allow measurement of DNL in the liver^12,16^. Enrichment of D2O in plasma was equal among the three main experimental groups (**Figure 1D**). GalNAc- siChREBP-treated rats exhibited a sharp decline in percent labeling of newly synthesized palmitate relative to either control group (**Figure 1E**). These findings are consistent with prior reports^7,10^, and with data in Figure 1C demonstrating reduced expression of key genes involved in DNL such as Fasn and Acly in response to ChREBP suppression. Interestingly, despite the decline in DNL, total palmitate levels were approximately doubled in livers from GalNAc-siChREPB-treated rats relative to either control group (**Figure 1F**). In addition, GalNAc-siChREBP treatment caused a marked increase in liver TG stores relative to either control group (**Figure 1G**), accompanied by a clear decrease in circulating TG (**Figure 1H**). The juxtaposition of increased hepatic TG with reductions in DNL and circulating TG is consistent with impaired hepatic VLDL packaging and secretion as noted in other liver ChREBP loss-of-function models^4,10,17,18^. This regulation of lipid export is thought to be mediated by transmembrane 6 superfamily 2 (Tm6sf2) and/or microsomal TG transfer protein (MTTP)^18–20^. Here we confirmed that GalNac-siChREBP treatment caused decreases in Tm6sf2 mRNA levels (**Figure 1C)** as well as Tm6sf2 protein levels as measured by immunoblot analysis (**Figure 1I**). GalNAc-siChREBP treatment also caused a significant increase in liver glycogen compared to either the GalNAc-siCtrl or saline-injected control groups (**Figure 1J**), as also found in glycogen storage diseases associated with downregulation of G6pc and Plkr (**Figure 1C**)^3,7,18,21^. Thus, knockdown of ChREBP caused the liver to increase storage of two major fuels, fatty acids in the form of TG and glucose in the form of glycogen. Consistent with these findings, GalNAc-siChREBP treatment caused a significant increase in liver weight relative to either control group (**Figure 1K**).

### Global transcriptomic effects of GalNAc-siChREBP treatment assessed by RNA-seq analysis

To gain insight into transcriptomic programs that may drive non-canonical actions of ChREBP, we performed RNA-seq analysis on liver from HF/HS-fed rats injected with GalNAc-siChREBP, GalNAc-siCtrl, or saline. Principal component analysis separated ChREBP knockdown from the two control groups (**Figure 2A**). However, the GalNAc-siCtrl control samples also separated from the saline control samples. Genes contributing to the discriminatory principal components included known regulators of lipid metabolism consistent with our qPCR findings (**Figure 1C** and **Supplemental Data 1**). Gene set enrichment analysis performed on differentially expressed genes from rats treated with GalNAc-siCtrl compared to saline identified genes involved in cholesterol and other sterol biosynthetic processes as well as genes involved in lipid metabolism and lipid droplet organization (**Supplemental Data 2**). These genes were enriched for targets of the transcription factor NR1I3, also known as Constitutive Androstane Receptor-beta (CAR-beta), which is known to be responsive to a range of xenobiotics including some sterols to regulate liver and systemic lipid metabolism^22^. Consistent with this, NR1I3/CAR-beta expression is reduced in livers from rats treated with GalNAc-siCtrl compared to saline (**Supplemental Data 2**). Thus, treatment of animals with GalNAc siRNA may have “off-target” effects on lipid metabolism mediated in part through effects on CAR-beta. These findings support a focused comparison of the GalNAc-siChREBP and GalNAc- siCtrl groups as the most appropriate and rigorous approach for defining ChREBP-specific effects on metabolism going forward, as it controls for non-specific effects of the GalNAc-mediated siRNA delivery method. Therefore, metabolites, transcripts or proteins that are different in the GalNAc- siChREBP-treated versus GalNAc-siCtrl-treated controls are emphasized in the data presentation and discussion that follows.

**Figure 2.**
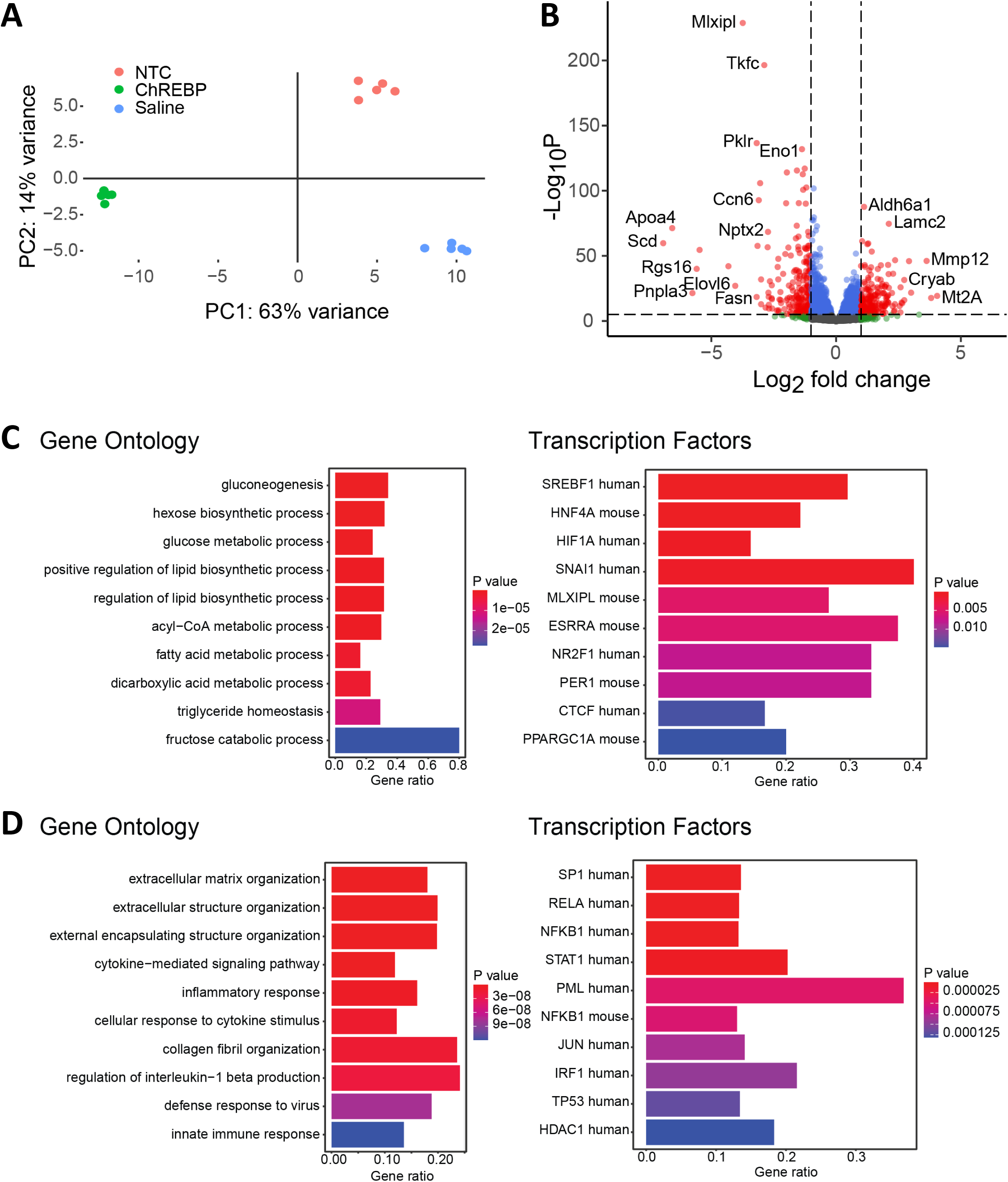
RNA-seq Analysis. RNA-seq was performed on liver samples obtained from rats fed HF/HS diet and treated with GalNAc-siChREBP, GalNAc-siCtrl, or saline over a 28 day period. **A.** Principal component plot for GalNAc-siCtrl- (red dots), GalNAc-siChREBP- (green dots), and saline- (blue dots) treated rats. Each dot represents an individual animal. **B.** Volcano plot depicting differentially expressed genes in livers from GalNAc-siChREBP versus GalNAc-siCtrl treated animals. **C.** Pathway analysis showing the top 10 gene ontology and transcription factor gene sets *downregulated* in GalNAc-siChREBP treated livers versus GalNAc-siCtrl. Color is indicative of p-value, and gene ratio corresponds to the fraction of genes in the gene set that contributed to enrichment. **D.** Pathway analysis showing the top 10 gene ontology and transcription factor gene sets *upregulated* in GalNAc-siChREBP treated livers versus GalNAc-siCtrl.

We further investigated differences between the GalNAc-siChREBP-treated and GalNAc-siCtrl-treated rats by “volcano plot” analysis of our RNA-seq data (**Figure 2B**). In liver samples from rats treated with GalNAc-siChREBP compared to treatment with GalNAc-siCtrl, ChREBP (Mlxipl) was the most significantly downregulated transcript, confirming the efficiency of our knockdown strategy (**Figure 2B and Supplemental Data 3**). Other well-known ChREBP target genes^1–11^ involved in glucose and fructose catabolism, glucose production, and lipogenesis were also significantly downregulated in response to GalNAc-siChREBP treatment (**Figure 1C and Supplemental Data 4)**. Gene set enrichment analysis demonstrated enrichment of down-regulated genes with known targets of ChREBP, as well as other key regulators of liver metabolism such as SREBF1, HNF4A and HIF1A (**Figures 2C and 2D).** We also found evidence for regulation of networks unrelated to core intermediary metabolic pathways. This included effects of ChREBP knockdown to increase expression of gene sets involved in extracellular matrix remodeling, inflammation, and immune responses **(Figures 2C and 2D).** Differentially regulated genes involved in these processes were enriched for targets of the transcription factors RELA and NFKB1, which are known to regulate stress and inflammatory responses. These results are consistent with a role for ChREBP in mediating the adaptive response to overnutrition^1,2^.

### Alterations in hepatic lipid metabolism in response to GalNac-siChREBP treatment

GalNAc- siChREBP treatment caused a decrease in circulating non-esterified fatty acids (NEFA) (**Figure 3A**), accompanied by an approximate 2.5-fold increase in circulating β-hydroxybutyrate and total ketone levels (**Figure 3B**). GalNAc-siChREBP treatment did not affect circulating glycerol levels (**Figure 3C**), indicating that adipose tissue lipolysis was not impaired. The fall in circulating NEFA, increases in liver TG and total palmitate, and increases in circulating ketones in the GalNAc-siChREBP group suggests increased hepatic fatty acid uptake and oxidation in response to hepatic ChREBP suppression. To further explore mechanisms underlying these changes, we used targeted tandem mass spectrometry to measure a panel of lipid-derived acylcarnitines in liver extracts. We found clear increases in a set of even chain acylcarnitines produced by fatty acid oxidation (FAO) in GalNAc-siChREBP treated rats compared to the two control groups (**Figure 3D**). This included intermediates produced in each round of the beta-oxidation spiral from palmitate or oleate, including C18:1, C16:0, C14:0, C14:1, C12:0, and C10:0 acylcarnitines, as well as a set of hydroxylated species of similar chain length thought to be generated by the individual FAO enzymes^23^. In addition, GalNAc-siChREBP treatment caused significant increases in β-hydroxybutyryl (C4-OH) carnitine, a marker of fatty acid oxidation to ketones^24^, as well as acetyl (C2) acylcarnitine.

**Figure 3.**
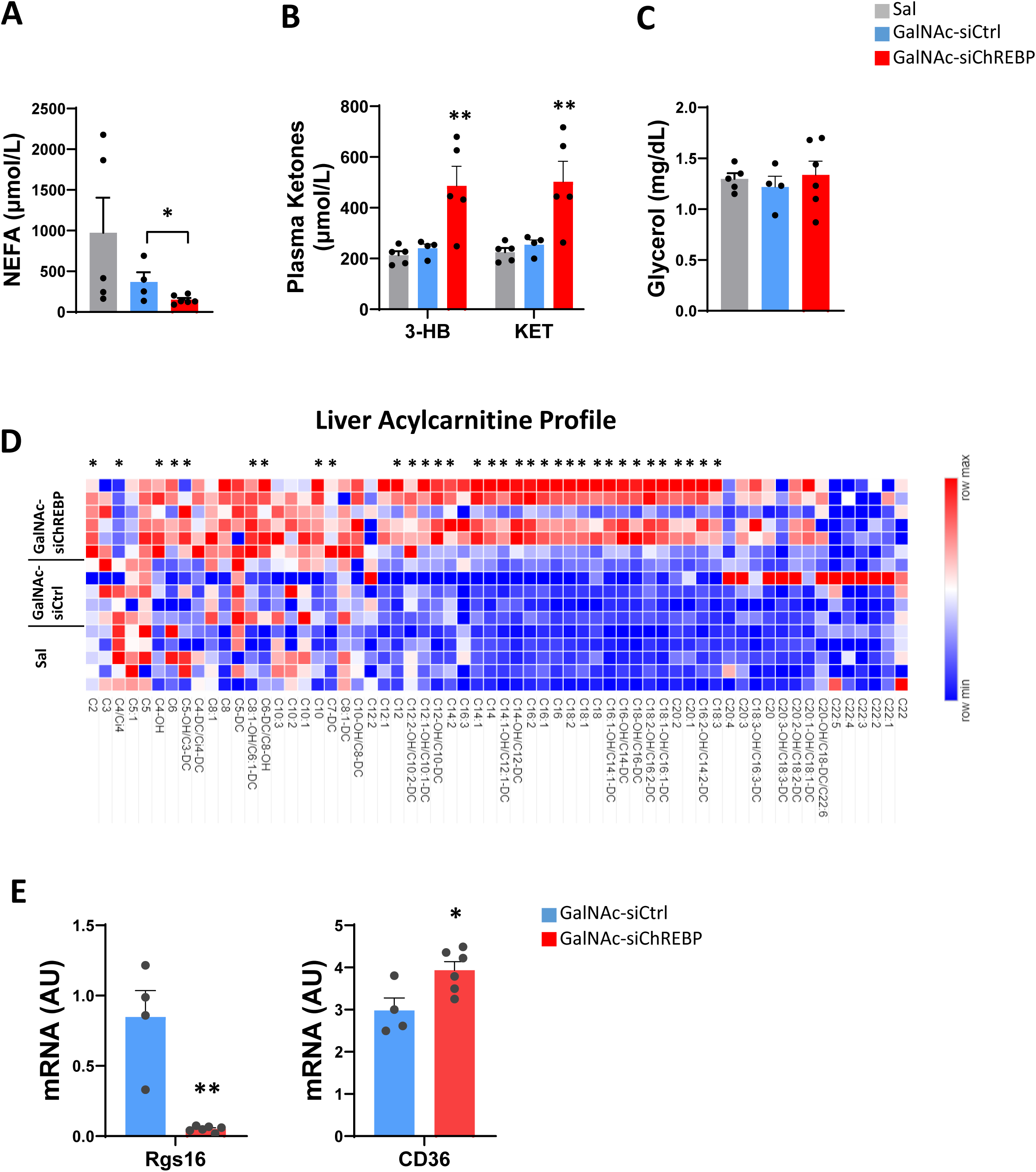
Effects of GalNAc-siRNA treatment on plasma and liver indices of lipid metabolism. Rats fed on HF/HS diet received multiple injections of GalNAc-siChREBP, GalNAc-siCtrl, or saline over a 28 day period prior to sacrifice and collection of plasma and liver samples for measurement of transcripts and metabolites reflective of changes in lipid metabolism. **A.** Plasma non-esterified fatty acids (NEFA) levels. **B.** Plasma β-hydroxybutyrate (3-HB) and total ketones (KET) levels. **C.** Plasma glycerol levels. **D.** Levels of individual acylcarnitines in liver samples measured by targeted tandem mass spectrometry. Heat map generated using Morpheus software (https://software.broadinstitute.org/morpheus). **E.** q-PCR analysis of Rgs16 (left), a G-protein suppressor of fatty acid oxidation, and CD36 (right), a fatty acid transporter in liver. Data are the mean ± standard error of the mean (SEM) for 6 independent rats/liver samples for the GalNAc- siChREBP-treated group and 5 rats/liver samples for the GalNAc-siCtrl and saline-treated groups. Statistical analysis performed using one-way ANOVA followed by Tukey test (A-D) or unpaired t-test (E). Symbols *, **, *** indicate a measure significantly different in the GalNAc-siChREBP treatment group compared to either the GalNAc-siCtrl or saline groups, with p < 0.05, p < 0.01 and p < 0.0001, respectively.

### RNA-seq identifies genes involved in altered lipid handling in response to ChREBP knockdown

In light of the foregoing changes in indices of fatty acid metabolism, we used our RNA-seq data set to identify changes in expression of transcripts encoding enzymes involved in fatty acid oxidation (FAO). Some, but not all enzymes of the mitochondrial FAO pathway were coordinately up-regulated in response to GalNAc-siChREBP treatment, including acyl CoA dehydrogenases ACAD11 (very long-chain; 1.4-fold increase, padj = 10^-10^) and ACADl (long chain; 1.2-fold increase, padj = 1 x 10^-6^), enoyl CoA hydratase Echs1 (1.2-fold increase, padj = 7.7 x 10^-5^), 3-hydroxyacyl CoA dehydrogenase HACD2 (1.1-fold increase, padj = 0.002) and beta-ketothiolase HADHB (1.1-fold increase, padj = 0.08), findings confirmed by q-PCR (**Supplemental Figure 2A**). Interestingly, medium chain ACAD expression did not change in response to ChREBP suppression, while expression of short chain ACAD (also known as SCHAD or Hadh) decreased sharply (1.6-fold decrease, padj = 2.9 x 10^-30^) (**Supplemental Figure 2A**). In contrast to these effects on transcripts encoding mitochondrial FAO enzymes, transcripts encoding core enzymes of peroxisomal fatty acid oxidation were significantly and coordinately downregulated in livers of GalNAc-siChREBP compared to GalNAc-siCtrl-treated rats, including acyl CoA oxidase-1 ACOX 1 (1.3-fold decrease, padj = 8.3 x 10^-11^), D-bifunctional protein D-BP (1.2-fold decrease, padj = 1.5 x 10^-8^) and both isoforms of acyl CoA acyltransferase 1 ACCA1 (ACCA1a, 1.8-fold decrease, padj = 2.9 x 10^-48^; ACCA1b, 1.9-fold decrease, padj = 1.3 x 10^-10^), findings confirmed by q-PCR (**Supplemental Figure 2A)**. Suppression of ChREBP expression caused sharp down-regulation of regulator of G protein 16 (Rgs16), a G-protein that suppresses FAO (**Figure 3F**)^25^. ChREBP suppression also increased expression of the CD36 fatty acid transporter (**Figure 3F**) and decreased levels of the Tm6sf2 transcript and protein (**Figure 1C and 1I**), potentially contributing to the increase in hepatic palmitate and TG levels as reported in **Figure 1**. Altogether, these transcriptomic changes suggest that ChREBP suppression activates influx of circulating fatty acids (via upregulation of CD36), to increase FAO (via downregulation of Rgs16 and induction of the core enzymes of mitochondrial fatty acid oxidation) resulting in increases in circulating ketones and hepatic fatty acid-derived acylcarnitines. We suggest that the increases observed in a wide set of fatty acid-derived acylcarnitine species reported in **Figure 3** could reflect increased entry of fatty acids into the β-oxidation spiral, but an inadequate capacity to fully oxidize the intermediates to CO2 in the TCA cycle, as has been previously described in skeletal muscles of rodents fed on high calorie diets^26^. This interpretation could be consistent with a prior study reporting suppression of FA oxidation in livers of mice with global ChREBP knockout^13^.

### Analysis of short-chain acyl CoA and free CoA

Seeking to expand our understanding of metabolic effects of ChREBP manipulation, we applied our previously described targeted metabolomics panel for coenzyme A (CoA) and CoA-modified metabolic intermediates^27^. We unexpectedly observed a striking decrease in free CoA levels in GalNAc-siChREBP-treated rats compared to GalNAc-siCtrl-treated controls (**Figure 4A**). This was accompanied by marked reductions in several short-chain acyl CoA metabolites, including acetyl CoA, butyryl CoA, HMG CoA, and propionyl CoA (**Figure 4B**).

**Figure 4.**
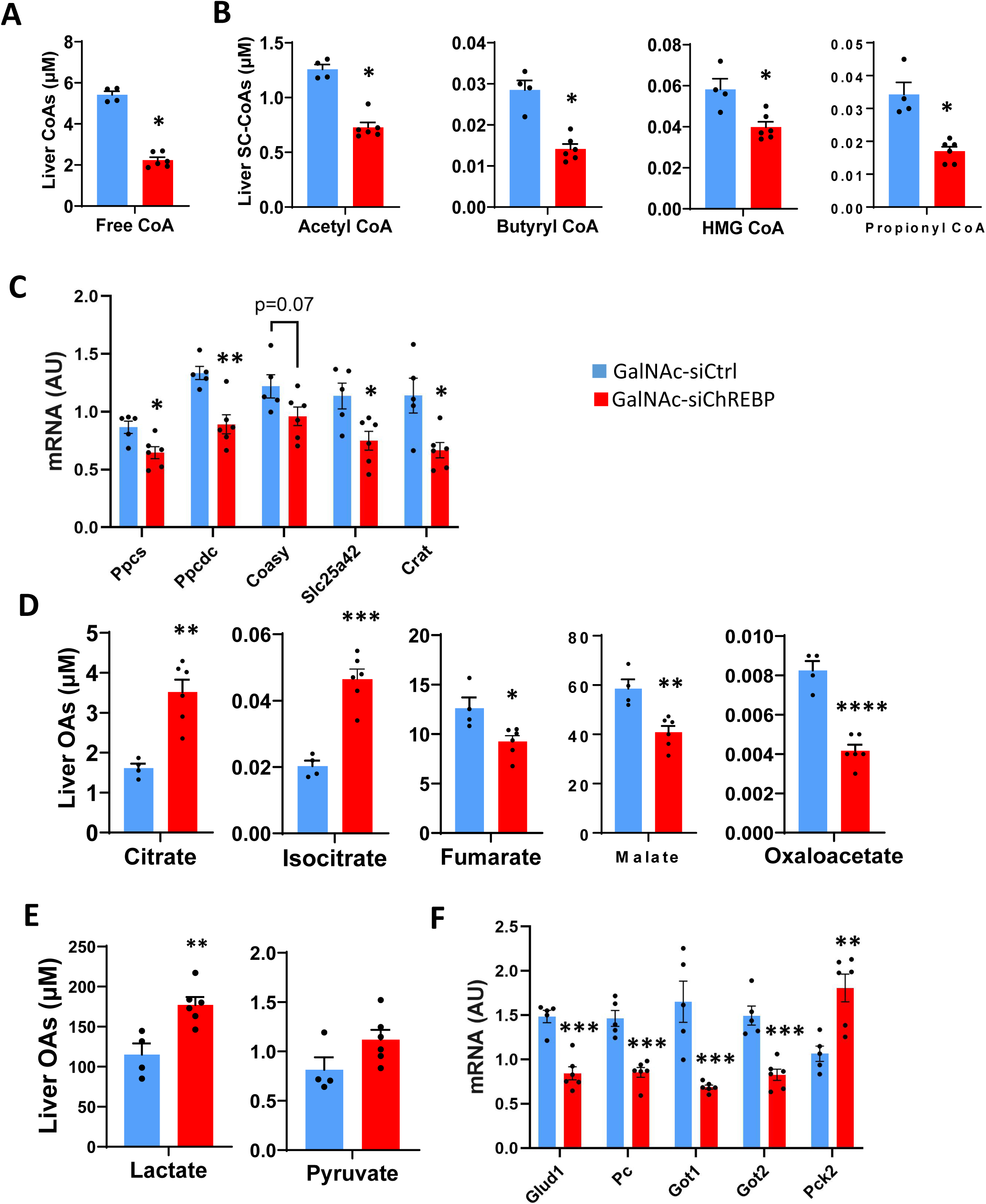
Effects of GalNAc-siChREBP treatment on indices of CoA, acyl CoA, and TCA cycle metabolism. Rats fed on HF/HS diet received multiple injections of GalNAc-siChREBP or GalNAc- siCtrl over a 28 day period. **A.** Free coenzyme A levels in liver. **B.** Levels of the short-chain acyl CoA intermediates acetyl CoA, butyryl CoA, HMG CoA, and propionyl CoA in liver. **C.** Expression of transcripts involved in metabolism of CoA and acyl CoAs measured by q-PCR, including phosphopantothenoylcysteine synthetase (Ppcs), phosphopantothenoylcysteine decarboxylase (Ppcdc), coenzyme A synthase (CoAsy), the mitochondrial CoA transporter (Slc25a), and carnitine O-acetyltransferase (Crat). **D.** Levels of TCA cycle intermediates in liver measured by targeted GC/MS. **E.** Lactate and pyruvate levels in liver. **F.** Expression of transcripts involved in supply of anaplerotic substrates to liver mitochondria measured by q-PCR, including glutamate dehydrogenase (Glud1), pyruvate carboxylase (Pc), both forms of aspartate transaminase (Got1 and Got2), and the mitochondrial isoform of PEPCK (Pck1). Data are the mean ± standard error of the mean (SEM) for 6 independent rats/liver samples for the GalNAc-siChREBP-treated group and 5 rats/liver samples for the GalNAc-siCtrl and saline-treated groups. Statistical analysis performed using unpaired t test. Symbols *, ** indicate a measure significantly different in the GalNAc-siChREBP treatment group compared to either the GalNAc-siCtrl group, with p < 0.05, <0.01, and <0.0001 respectively.

These findings led us to query our RNA-seq data set for genes involved in CoA biosynthesis and metabolism of short-chain acyl CoA metabolites. Three transcripts encoding enzymes of CoA synthesis from pantothenic acid were significantly reduced by GalNAc-siChREBP treatment— phosphopantothenoylcysteine synthetase (PPCS; 1.4-fold decrease, padj = 5.5 x 10^-4^), phosphopantothenoylcysteine decarboxylase (PPCDC; 1.4-fold decrease, padj = 6.6 x 10^-5^), and coenzyme A synthase (CoAsy; 1.3-fold decrease, padj = 5.3 x 10^-13^), and these changes were confirmed by q-PCR **(Figure 4C**). GalNAc-siChREBP treatment also resulted in reduced expression of the transcript encoding the mitochondrial CoA transporter (Slc25a42, 1.4-fold decrease, padj = 4.4 x 10^-10^), also confirmed by q-PCR **(Figure 4C**). In addition, the transcript encoding carnitine O-acetyltransferase (CRAT) was decreased by GalNAc-siChREBP treatment (1.9-fold decrease, padj = 0.00049) and confirmed by RT-PCR (**Figure 4C**). CRAT catalyzes the reversible conversion of free CoA + short-chain acylcarnitines to short-chain acyl CoAs + carnitine. We hypothesize that the decreases observed in multiple short-chain acyl CoA metabolites (**Figure 4B**) may be driven by the strong decrease in the free CoA pool and the reduction in CRAT activity. These factors may also explain how acetyl CoA levels fall (**Figure 4B**) while acetylcarnitine levels increase (**Figure 3D**) in GalNAc-siChREBP relative to GalNAc-Ctrl-treated rats, as conversion of acetyl carnitine to acetyl CoA would be impaired by low CoA levels and reduced CRAT activity.

### Analysis of TCA cycle intermediates

Our finding of reduced levels of CoA and expression of CoA synthetic enzymes point to previously unanticipated changes in mitochondrial fuel metabolism resulting from ChREBP knockdown. Burgess, et al previously reported that global ChREBP knockout in mice results in reduced hepatic glucose catabolism, but increased lactate and pyruvate oxidation to maintain energy balance in the face of impaired FA oxidation^13^. They also reported an increase in pyruvate dehydrogenase enzymatic activity without a change in PDH protein amount, and a fall in expression of the PDK3 isoform, an inhibitory PDH kinase^13^. Here we found either a modest decrease or no change in expression of transcripts encoding a subset of PDH component subunits (PDHA1, PDHX, and DLAT), with a decrease in the level of the transcript encoding PDHB (reduced 2.0-fold, padj = 2.6 x 10^-14^), and the predominant PDH kinase PDK4 (reduced 2-fold, padj = 8.7 x 10^-8^) with no significant change in expression of PDK3, findings still consistent with an overall increase in hepatic PDH activity (**Supplemental Figure 2B**). To gain further insight into changes in mitochondrial function in response to ChREBP knockdown, we used targeted GC/MS to measure a panel of organic acids/TCA cycle intermediates by a previously described method^28^. We found increases in hepatic citrate and isocitrate levels in GalNAc-siChREBP compared to GalNAc-siCtrl-treated rats, coupled with decreased levels of the more distal TCA cycle intermediates fumarate, malate, and oxaloacetate (**Figure 4D**). Pyruvate levels showed a trend to increase in the GalNAc-siChREPB-treated rats whereas lactate levels were significantly increased (**Figure 4E**).

The fall in oxaloacetate and acetyl CoA levels and increase in citrate and isocitrate that we report in **Figure 4** could be consistent with an increase in acetyl CoA and oxaloacetate consumption through the PDH, citrate synthase and isocitrate dehydrogenase reactions to form citrate and isocitrate. Another possible contributor to the rise in the citrate and isocitrate pools in livers of GalNAc-siChREBP-treated rats could be the decrease in ACLY expression that occurs in response to ChREBP suppression **(Figure 1C**), limiting the use of citrate to form acetyl CoA as a substrate for DNL. This framework raises the question of the source of pyruvate for driving PDH flux, especially in light of the sharp decrease in pyruvate kinase expression (**Figure 1C**). To investigate this, we examined the expression of the family of monocarboxylic acid carriers, including 4 isoforms known to be pyruvate and lactate transporters, and found increases in transcript levels for 3 of the 4--SLC16a3, SLC16a4, and SLC16a7 (1.2-fold increase, padj = 0.0004; 1.2-fold increase, padj = 0.004; 1.7-fold increase, padj = 0.009, respectively) with no significant change in expression of the fourth isoform SLC16a1 (**Supplemental Figure 2C**). Among members of the SLC16a family of transporters with specificity for other substrates, a transporter of aromatic amino acids (SLC16a10) was significantly increased, whereas transporters for GABA and thyroid hormone T3 (SLC16a11 and SLC16a2) were significantly decreased, and several other family members exhibited no significant change in expression (SLC16a6, SLC16a9, SLC16a12, SLC16a13) (**Supplemental Figure 2D**), highlighting the concerted and selective pattern of increased expression of pyruvate/lactate transporters in our data set. Interestingly, the transcripts for lactate dehydrogenase a (LDHa) and glutamate/pyruvate transaminase 2 (GPT2) were both decreased in liver in response to GalNAc-siChREBP treatment (2.3-fold decrease, p = 1.07 x 10^-31^ and 1.5-fold decrease, p = 6.4 x 10^-5^, respectively), as confirmed by q-PCR (**Supplemental Figure 2E**), possibly serving as a mechanism for preserving the pyruvate pool for oxidation via the PDH reaction.

Turning to mitochondrial pathways, a surprising finding is that the clear increases in citrate and isocitrate levels in livers of GalNAc-siChREBP-treated rats occurred despite strong induction of 2- ketoglutarate dehydrogenase expression (OGDH, the enzyme that converts citrate to succinyl CoA; 0.52-fold increase, padj = 6.9 x 10^-24^). We interpret this to mean that the marked depletion of free CoA in GalNAc-siChREBP-treated rats limits its supply as a cofactor for the OGDH reaction, contributing to accumulation of the proximal metabolites citrate and isocitrate even in the face of compensatory induction of the OGDH gene, accompanied by decreased levels of the more distal TCA cycle metabolites succinate, fumarate and malate.

These data prompted examination of the expression of key anaplerotic enzymes that feed carbon into the TCA cycle. All those examined were found to be strongly downregulated in livers from GalNAc-siChREBP-treated compared to GalNAc-siCtrl-treated rats, including glutamate dehydrogenase (GLUD1, padj = 1.34 x 10^-15^), pyruvate carboxylase (PC, padj = 2.99 x 10^-21^), and both forms of aspartate transaminase (GOT1, padj = 1.5 x 10^-12^, and GOT2, padj = 1.6 x 10^-109^), all confirmed by q-PCR (**Figure 4F**). This raises the question of how the livers of GalNAc-siChREBP- treated rats are able to generate anaplerotic substrates for formation and expansion of the citrate and isocitrate pools as shown in **Figure 4D**. One possibility is the mitochondrial isoform of PEPCK (PCK2), which normally generates phosphoenolpyruvate (PEP) from oxaloacetate and GTP in an irreversible reaction, but which has been shown to function as an anaplerotic enzyme that catalyzes oxaloacetate formation from PEP in bacteria where pyruvate kinase is knocked out^29^. Here, pyruvate kinase expression was essentially completely silenced by GalNAc-siChREBP treatment (**Figure 1C**), perhaps helping to explain the clear, possibly compensatory induction of PCK2 (2.0-fold increase, p = 1.4 x 10^-33^), as confirmed by q-PCR (**Figure 4F**).

### Effects of ChREBP on amino acid metabolism

Targeted profiling of hepatic amino acid levels by tandem mass spectrometry revealed significant decreases in hepatic Gly, Leu/Ile (a combined measure of leucine and isoleucine), Asx (a combined measure of aspartate and asparagine), ornithine, and Tyr levels in livers of GalNAc-siChREBP compared to GalNAc-siCtrl-treated animals (**Figure 5A**). In addition, RNA-seq analysis revealed induction of multiple transcripts encoding plasma membrane amino acid transporters in response to ChREBP knockdown. Transcripts significantly increased in response to GalNAc-siChREBP compared to GalNAc-siCtrl treatments included the large neutral amino acid/BCAA carrier (LAT1, Slc7a5), the alanine/taurine carrier (Slc6a6), the glutamate transporter (Slc1a2), the neutral amino acid transporter for Thr, Ser, Cys, Ala, and Gln (ASCT2, Slc1a5), the arginine/histidine transporter (Slc7a1), and the glutamine transporter (Slc38a1), all confirmed by qPCR **(Figure 5B**). Of note, the transcript encoding the sodium-coupled neutral amino acid transporter-2 (SNAT2, Slc38a2), previously reported by another group to be suppressed by ChREBP^30^, was not significantly altered by GalNAc-siChREBP treatment in our studies (**Figure 5B**). Overall, the observed increases in amino acid transporter expression, coupled with the decreases in levels of multiple liver amino acids, suggests that ChREBP suppression may activate amino acid catabolism as an alternative energy source, leading to amino acid depletion accompanied by a compensatory upregulation of hepatic amino acid transporters to offset this decline.

**Figure 5.**
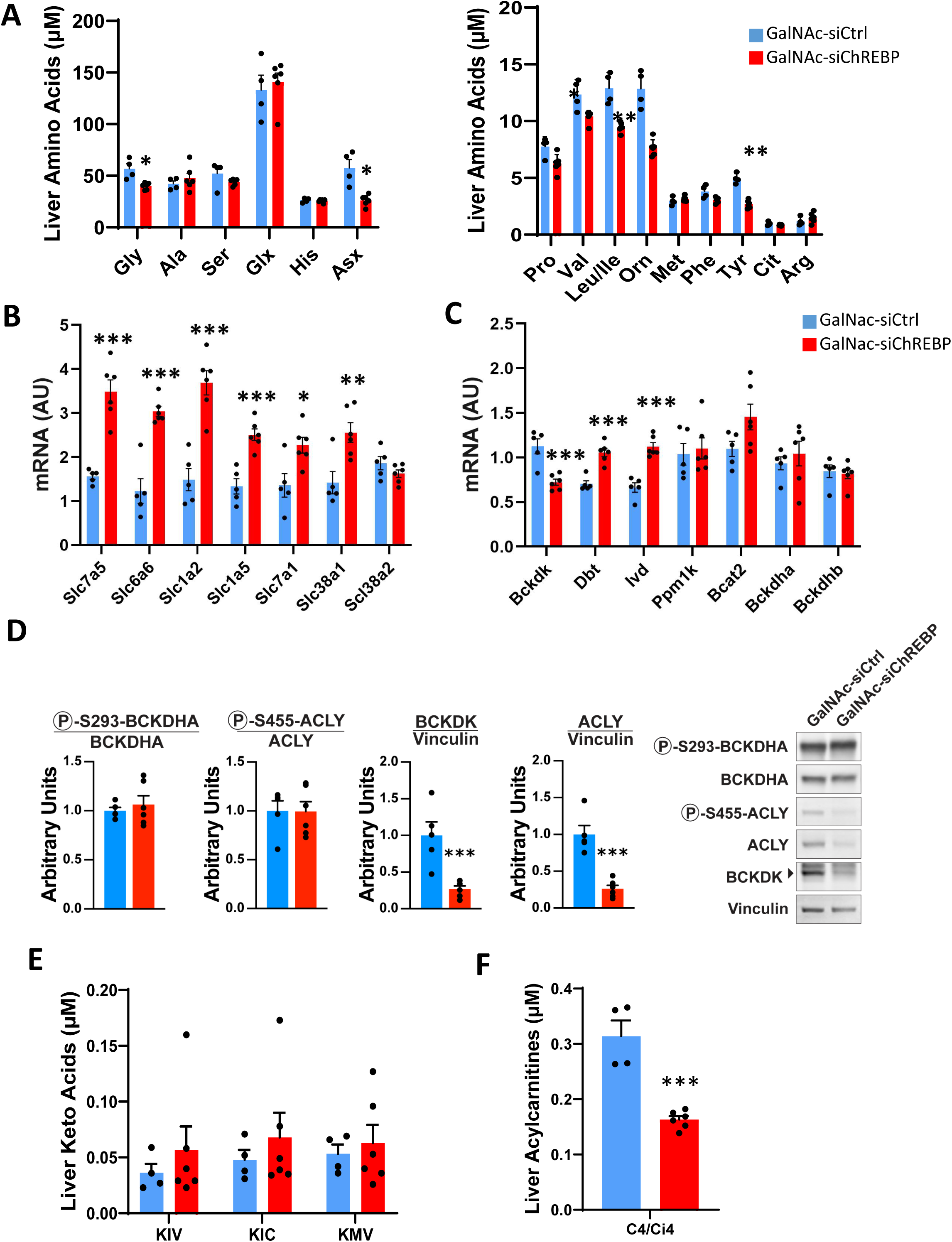
Effects of GalNAc-siChREBP on indices of amino acid metabolism. Rats fed on HF/HS diet received multiple injections of GalNAc-siChREBP or GalNAc-siCtrl over a 28 day period. **A.** Liver amino acid levels measured by targeted tandem mass spectrometry. **B.** q-PCR analysis of transcripts encoding amino acid transporters identified as differentially regulated by RNA-seq analysis. Note that Slc38a2 (SNAT2) is the only transcript shown in the this panel that was not significantly regulated by GalNAc-siChREBP treatment, consistent with the RNA-seq data. **C.** Regulation of transcripts encoding proteins involved in BCAA catabolism, including branched-chain ketoacid dehydrogenase kinase (Bckdk), lipoamide acyltransferase (Dbt), isovaleryl CoA dehydrogenase (Ivd), the BCKDH phosphatase (PPM1K), branched-chain aminotransferase-2 (Bcat2), branched-chain ketoacid dehydrogenase alpha subunit (Bckdha) and branched-chain ketoacid dehydrogenase beta subunit (Bckdhb). **D.** Representative immunoblot (right) and densitometric quantification (left) of phosphorylated (P-S293-BCKDHA) and total (BCKDHA) branched-chain ketoacid dehydrogenase A, phosphorylated (P-S455-ACLY) and total (ACLY) ATP-citrate lyase, branched-chain ketoacid dehydrogenase kinase (BCKDK), and vinculin as a loading control. **E.** Levels of the branched-chain ketoacids ketoisovalerate (KIV), ketoisocaproate (KIC) and ketomethylvalerate (KMV) measured by targeted LC-MS/MS in liver. **C.** Levels of 2- hydroxybutyrate (2-HB) in liver measured by targeted LC-MS/MS. **D.** Levels of C4/Ci4 acylcarnitine in liver measured by targeted tandem mass spectrometry. Data are the mean ± standard error of the mean (SEM) for 6 independent rats/liver samples for the GalNAc-siChREBP-treated group and 5 rats/liver samples for the GalNAc-siCtrl and saline-treated groups. Statistical analysis performed using unpaired t-test or multiple t-test. Symbols *, **, *** indicates a measure significantly different in the GalNAc-siChREBP treatment group compared to either the GalNAc-siCtrl group, with p < 0.05, p < 0.01 and p < 0.005, respectively.

We also examined the impact of GalNAc-siChREBP treatment on expression of a broad set of genes involved in BCAA catabolism. In agreement with our recent finding that overexpression of ChREBP-β increases expression of BCKDK in liver^12^, administration of GalNAc-siChREBP decreased BCKDK mRNA levels, which would be expected to result in increased BCKDH activity and increased BCAA catabolism (**Figure 5C**). Consistent with this model, ChREBP knockdown also caused an increase in expression of the E2 component (lipoamide acyltransferase, Dbt) of the BCKDH complex, as well as an increase in the transcript encoding isovaleryl CoA dehydrogenase (Ivd), the third enzyme in catabolism of leucine (**Figure 5C**). Other genes involved in BCAA metabolism including Ppm1K, Bcat2, Bckdha, and Bckdhb were not significantly affected by ChREBP knockdown, although a trend for increased expression of the BCAT2 transcript (p = 0.07) was observed in response to GalNAc- siChREBP treatment (**Figure 5C**). Treatment with GalNAc-siChREBP caused a 75% decrease in BCKDK protein levels relative to GalNAc-siCtrl-treated rats, but had no effect on BCKDH phosphorylation (**Figure 5D**). Also, GalNAc-siChREBP treatment reduced phospho-S455 ACLY levels relative to both controls, but this effect could be accounted for by a reduction in total ACLY protein (**Figure 5D**).

As previously noted, we found that GalNAc-siChREBP treatment tended to cause a decrease in liver valine (p = 0.052) and caused a significant decrease in leucine/isoleucine (p = 0.002) relative to GalNAc-siCtl treatment (**Figure 5A**). However, GalNAc-siChREBP treatment had no effect on levels of the branched-chain keto acids, ketoisovalerate (KIV), ketoisocaproate (KIC), or ketomethylvalerate (KMV) (**Figure 5E**). GalNAc-siChREBP treatment also decreased the level of the valine-derived metabolite butyryl/isobutyryl (C4/Ci4) acylcarnitine compared to GalNAc-siBCKDK treatment (**Figures 5F**). These data suggest that GalNAc-siChREBP treatment activates consumption of BCAA but not the BCKA, while causing a decrease in a valine-derived metabolite relative to GalNAc- siBCKDK treatment. These data, and the absence of any changes in BCKDH or ACLY phosphorylation despite the 75% reduction in BCKDK protein levels, suggest that ChREBP exerts regulatory influences on BCKDH activity and BCAA metabolism beyond the regulation afforded by BCKDK and PPM1K-mediated phosphorylation, a topic that remains to be explored in future studies.

### Effects of ChREBP on nucleotide metabolism

Given the multiple unexpected effects of ChREBP suppression on diverse aspects of cellular metabolism, including pathways expected to influence cellular energy homeostasis, we next assessed the effects of ChREBP suppression on nucleotide and purine pathway intermediates using our previously described targeted LC-MS method for their measurement^31^. We found levels of PRPP, ZMP, and AICAR to be dramatically increased in livers of GalNAc-siChREBP-treated relative to GalNAc-siCtrl-treated rats (**Figures 6A and 6B**). PRPP, ZMP, and AICAR are late intermediates in the *de novo* purine biosynthesis pathway, and PRPP is also a co-substrate in the last steps of the *de novo* NAD synthesis pathway and purine salvage pathways. In contrast to the increase in these precursors, the products of the purine synthesis pathways, IMP, GMP and AMP, as well as products of the *de novo* nicotinamide dinucleotide synthesis pathway, including nicotinic acid mononucleotide (NaMN), NAD(P) and NAD(P)H, were all sharply reduced in livers of GalNAc-siChREBP compared to GalNAc-siCtrl-treated rats (**Figures 6A, 6B, and 6C**). Interestingly, levels of the nucleotide diphosphates (ADP, UDP, GDP) or nucleotide triphosphates (ATP, UTP, GTP) were not lowered by GalNAc-siChREBP treatment, and UTP was significantly increased (**Figure 6D)**.

**Figure 6.**
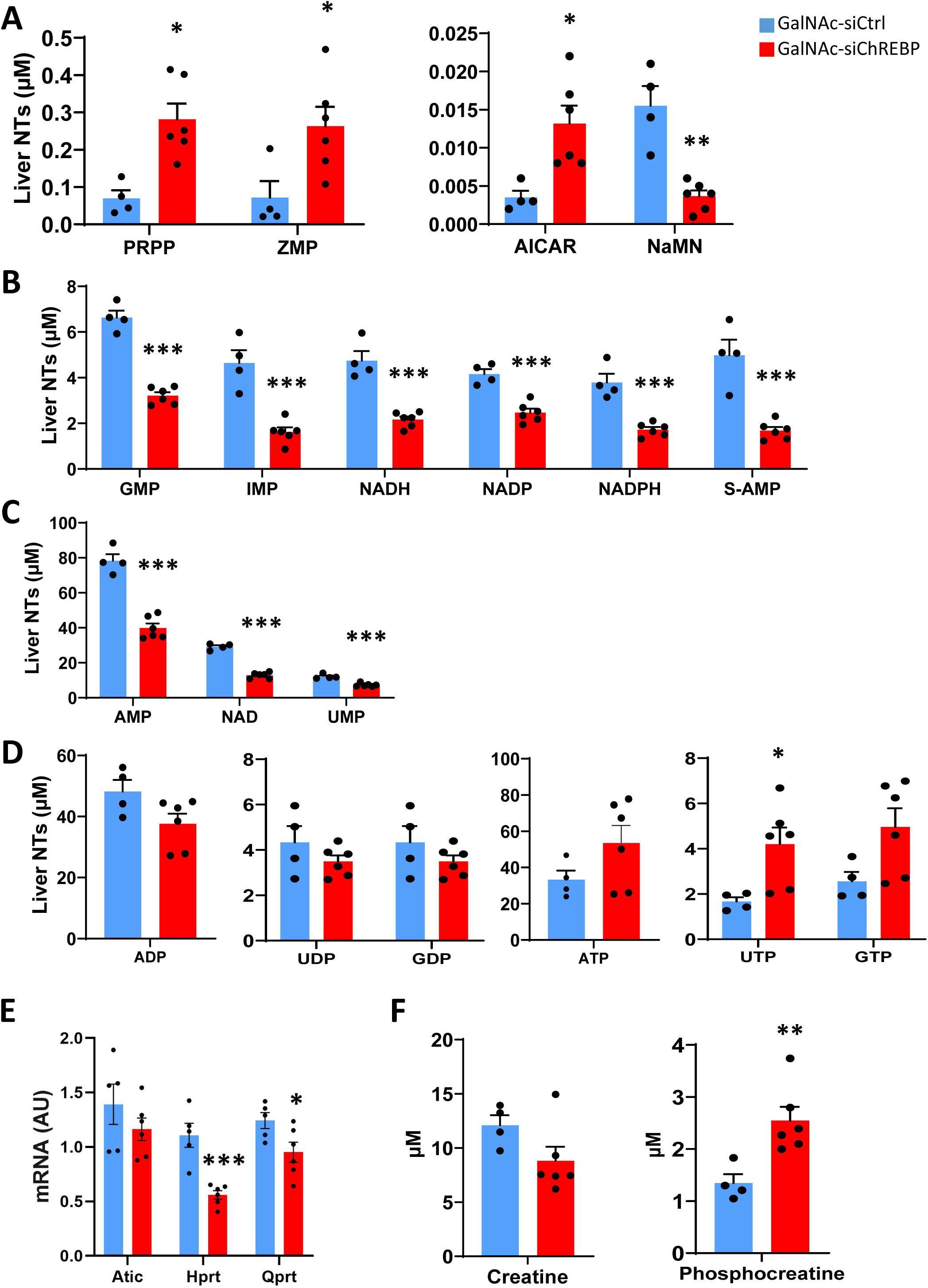
Effects of GalNAc-siChREBP on indices of nucleotide metabolism in liver. Rats fed on HF/HS diet received multiple injections of GalNAc-siChREBP or GalNAc-siCtrl over a 28 day period prior to sacrifice and analysis of transcripts and metabolites related to nucleotide metabolism. **A.** Levels of purine and nicotinamide dinucleotide pathway intermediates PRPP, ZMP, AICAR and NaMN measured by targeted LC/MS. **B.** Levels of nucleotide and nicotinamide dinucleotide pathway intermediates GMP, IMP, NADH, NADP, NADPH, and S-AMP measured by targeted LC/MS. **C.** Levels of nucleotide and nicotinamide dinucleotide pathway intermediates including AMP, NAD, and UMP measured by targeted LC/MS. **D.** Levels of nucleotide di- and tri-phosphate intermediates, including ADP, UDP, GDP, ATP, UTP, and GTP measured by targeted LC/MS. **E.** Levels of key transcripts involved in *de novo* or salvage pathways of purine or nicotinamide dinucleotide synthesis in liver measured by q-PCR, including 5-aminoimidazole-4-carboxamide ribonucleotide formyltransferase/inosine monophosphate cyclohydrolase (Atic), hypoxanthine guanine phosphoribosyltransferase (Hprt), and quinolinate phosphoribosyltransferase (Qprt). **F.** Levels of creatine and phosphocreatine in liver. Data are the mean ± standard error of the mean (SEM) for 6 independent rats/liver samples for the GalNAc-siChREBP-treated group and 5 rats/liver samples for the GalNAc-siCtrl and group. Statistical analysis performed using unpaired t-test or multiple t-test. Symbols *, **, *** indicate a measure significantly different in the GalNAc-siChREBP treatment group compared to the GalNAc-siCtrl group, with p < 0.05, p < 0.01 and p < 0.005, respectively.

To further investigate mechanisms underlying the coupling of increases in levels of intermediates in the purine and nicotinamide synthesis pathways such as PRPP and AICAR with decreased levels of the end product nucleotide monophosphates (AMP, IMP, GMP) and nicotinamide dinucleotide pathway intermediates and products NaMN, (NAD(P), and NAD(P)H), we queried our RNA-seq data set for genes involved in the *de novo* biosynthesis and salvage pathways for these important mediators of energy homeostasis. The 5-aminoimidazole-4-carboxamide ribonucleotide formyltransferase/inosine monophosphate cyclohydrolase (ATIC) gene encodes a bifunctional enzyme that includes AICAR transformylase and IMP cyclohydrolase activities to catalyze conversion of AICAR to IMP with N^10^-formyl tetrahydrofolate as co-substrate for the reaction. ATIC transcript levels were significantly downregulated in livers of GalNAc-siChREBP- treated compared to GalNAc-siCtrl-treated rats by RNA-seq analysis (1.3-fold decrease, padj =.0001), and trended to decrease when measured by RT-PCR (**Figure 6E**). An alternate source of nucleotide monophosphates is the salvage pathway, with hypoxanthine guanine phosphoribosyltransferase (HPRT) serving as a final step in the pathway via its conversion of hypoxanthine to IMP, using PRPP as a co-substrate. HPRT transcript levels are were sharply decreased in GalNAc-siChREBP-treated compared to GalNAc-siCtrl-treated rats measured both by RNA-seq analysis (1.9-fold decrease, padj = 3.7 x 10^-32^), and q-PCR (**Figure 6E**). With regard to the changes in nicotinamide dinucleotide pools, the penultimate enzyme in the *de novo* biosynthesis pathway is quinolinate phosphoribosyltransferase (QPRT) which converts quinolinate derived from the tryptophan/kynurenine metabolism to NaNM and NAD, again using PRPP as a co-substrate. QPRT transcript levels were significantly reduced in GalNAc-siChREBP compared to GalNAc-siCtrl-treated rats both in the RNA-seq analysis (1.3-fold reduced, padj = 5.5 x 10^-11^) and when measured by q-PCR (**Figure 6E**), consistent with the increased levels of PRPP and lowered levels of NaNM, NAD(H), and NADP(H) in GalNAc-siChREBP-treated rats. These concerted effects of knockdown of ChREBP to suppress synthesis of the nucleotide monophosphate and nicotinamide dinucleotide pools surprisingly does not manifest in a decrease in the nucleotide diphosphate or nucleotide triphosphate pools (**Figure 6D**). One possible contributor to maintenance of these pools is the putative increase in pyruvate oxidation as described earlier, suggesting that oxidation of circulating pyruvate/lactate is sufficient to maintain hepatic energy balance. We also measured intermediates in phosphocreatine metabolism and found levels of phosphocreatine to be significantly increased, while levels of creatine exhibited a non-significant trend for decrease in livers of GalNAc-siChREBP-treated compared to GalNAc-siCtrl treated rats (**Figure 6F**). Taken together, these data unveil unanticipated roles of ChREBP in control of nucleotide metabolism and energy homeostasis.

## Discussion

In this study, we have integrated an extensive set of transcriptomic and metabolomic analyses in an effort to better understand the role of hepatic ChREBP in coordinated regulation of key pathways of intermediary metabolism. Since its discovery in 2001^32^, ChREBP has emerged as a central factor in regulation of tissue responses to carbohydrate ingestion^1,33^. It is particularly responsive to fructose, which is consumed in high amounts in modern society via sugar-sweetened beverages containing sucrose or high-fructose corn syrup. Consumption of fructose-containing foods and beverages produces carbohydrate metabolites that allosterically activate ChREBP-α, which acts upon an alternative promoter in the ChREBP gene to stimulate transcription of ChREBP-β, the more potent transcriptional activator among the two ChREBP isoforms^34^. The transcriptional programs activated by ChREBP-α and ChREBP-β enhance flux of carbohydrates to form lipids via gene products in the glycolytic and DNL pathways such as PKLR, ACLY, and FASN. ChREBP also has homeostatic functions in response to carbohydrate feeding, such as activation of glucose-6-phosphatase expression to limit accumulation of glucose-6-phosphate and glycogen during periods of increased carbohydrate influx^3,18,21^, and activation of transmembrane 6 superfamily member 2 (TM6SF2) expression^18^ to stimulate lipid export in VLDL particles, thereby limiting TG overaccumulation in the liver.

Here, via integration of targeted metabolomics and transcriptomics analyses, we have uncovered heretofore unrecognized functions of ChREBP in regulation of metabolic pathways that are associated with, but distinct from, core pathways of glucose and lipid metabolism. Our approach to data analysis was to allow ChREBP-mediated metabolite changes to point us to pathways in which to query potential underlying transcriptomic changes. This method identified pathways such as CoA synthesis, short-chain acyl CoA metabolism, and purine nucleotide and nicotinamide adenine dinucleotide synthesis that were not identified by cluster or pathway analyses of the transcriptomic data, suggesting an alternative metabolite-driven approach to integration of multi-omic data sets. By focusing on transcripts and metabolites specifically altered by GalNAc-siChREBP versus GalNAc-siCtrl treatment of HF/HS-fed rats, pathways identified as regulated by ChREBP included: 1) Biosynthesis of free coenzyme A (CoA), represented by decreased expression of 4 genes involved in CoA biosynthesis and transport, accompanied by a sharp lowering of hepatic free CoA levels. Several short-chain acyl CoA metabolites were also decreased, including acetyl CoA, butyryl CoA, HMG CoA, and propionyl CoA, presumably secondary to the depletion of the free CoA pool; 2) Regulation of a broad array of genes involved in pyruvate metabolism, including multiple isoforms of the monocarboxylic acid carrier (SLCa16) gene family, LDHa, glutamate/pyruvate transaminase 2 (GPT2), PKLR, and PDK4, which in aggregate may “funnel” pyruvate for use in PDH-mediated oxidation. Consistent with these changes in gene expression, we found decreased levels of acetyl CoA and oxaloacetate coupled with strong increases in citrate and isocitrate levels, suggesting increased flux through the citrate synthase reaction downstream of PDH; 3) *De novo* and salvage pathways of purine biosynthesis were suppressed. A block in these pathways was highlighted by the large increase in the co-substrates PRPP and AICAR and the fall in nucleotide monophosphate levels (AMP, IMP, GMP), aligned with suppression of transcripts encoding the key *de novo* pathway enzyme ATIC and salvage pathway enzyme HPRT. Nicotinamide mononucleotide (NaNM) and nicotinamide adenine nucleotides (NADP(H) and NAD(H)) were also decreased, aligned with lowered expression of the transcript encoding QPRT, a late step in the *de novo* synthesis pathway and another PRPP utilizing enzyme; 4) Suppression of transcripts encoding multiple anaplerotic enzymes, including PC, GDH, GOT1, and GOT2, coupled with an increase in expression of PCK2, the mitochondrial isoform of PEPCK, which may provide an alternative source of oxaloacetate; 5) Effects on amino acid metabolism, including lowering of Gly, Leu/Ile, ornithine, and Tyr levels, accompanied by upregulation of a set of transcripts encoding plasma membrane amino acid transporters.

These broad effects of ChREBP on diverse metabolic pathways have been revealed by suppressing expression of the transcription factor in a liver-selective fashion with GalNAc-siRNA technology, in rats fed on HF/HS diet. These findings predict the inverse effects of ChREBP under conditions where its activity is induced, such as in response to diets containing sucrose or fructose. In this context, the predicted concerted induction of genes involved in CoA synthesis can be interpreted as a function of ChREBP that allows conversion of the carbohydrate substrates fructose and glucose to CoA-modified intermediates required for mitochondrial energy metabolism and DNL, as well as modification of newly synthesized fatty acids to form fatty acyl CoAs, a required step for their utilization in both esterification and oxidation reactions. Similarly, ChREBP-mediated activation of glucose and fructose metabolism via well-known targets such the GLUT-2 and GLUT-5 glucose transporters, glucokinase regulatory protein, ketohexokinase, and pyruvate kinase must be accompanied by expression of anaplerotic enzymes to fuel increased TCA cycle flux for generation of cataplerotic/lipogenic intermediates, consistent with our finding of robust regulation of four such enzymes—PC, GDH, GOT1, and GOT2. Interestingly, the gene encoding the mitochondrial isoform of PEPCK, PCK2, is regulated in an opposite fashion to these four genes, suggesting that its suppression by ChREBP may contribute to conservation of TCA substrates via preventing PCK2-mediated conversion of oxaloacetate to PEP in the mitochondria. Our study also expands upon and helps explain earlier observations of increased pyruvate utilization in liver of global ChREBP knockout in mice^13^ via our new observations of upregulation of multiple monocarboxylic acid carrier family members to facilitate pyruvate uptake, preservation of the pyruvate pool by suppression of LDHa and GPT2, and activation of PDH flux by lowering of the most relevant PDK family member PDK-4. Interestingly, in the earlier study, feeding of mice with global knockout of ChREBP on a high sucrose diet resulted in starvation and death, suggesting that a switch to high reliance on pyruvate as the major oxidative fuel becomes untenable in the global knockout setting, whereas in our studies of liver-selective ChREBP knockdown, body weight and survival can be maintained even with feeding of HF/HS diet.

Another novel finding of our study is the effect of ChREBP to regulate key distal enzymes in both the *de novo* and salvage pathways for purine synthesis, as well as the *de novo* pathway of nicotinamide dinucleotide synthesis. Interestingly, a key intermediate in these pathways is PRPP, which is generated as a byproduct of glucose metabolism through the pentose monophosphate shunt. The increase in this glucose and fructose-derived metabolite is surprising given our finding of dramatic decreases in the transcripts encoding the GLUT-2 glucose transporter SLC2a2 and the GLUT-5 fructose transporter SLC2a5. Our observation of a clear increase in PRPP levels in rats treated with GalNAc-siChREBP therefore seems best explained by reliance on an alternate source of ribose-5- phosphate for PRPP synthesis, possibly nucleotide recycling. Placing this finding in the context of the known activation of ChREBP by carbohydrate feeding suggests that disposal of glucose and fructose requires coordinated upregulation of nucleotide and nicotinamide dinucleotide synthesis to support energy and redox-requiring pathways that mediate hexose disposal, including glycogen synthesis, DNL, acylation of fatty acids, lipid esterification and packaging, and possibly other cellular responses such as increased proliferation in the setting where anaplerotic/biosynthetic substrate is abundant. Whether the observed regulation of the ATIC, HPRT, and QPRT genes involved in nucleotide and nicotinamide dinucleotide synthesis is a direct effect of ChREBP or a result of an indirect action(s) remains to be determined.

Another novel finding of our study is the effect of ChREBP suppression to increase expression of a set of plasma membrane amino acid carriers, including LAT1/Slc7a5, Slc6a6, Slc1a2, ASCT2/Slc1a5, Slc7a1, and Slc38a1. All were identified as upregulated in our RNA-seq analysis, findings then confirmed by targeted q-PCR measurements. These changes in gene expression were accompanied by decreases in the levels of multiple amino acids in liver, suggesting that upregulation of the transporters may serve to compensate for increased rates of amino acid utilization. Our data are consistent with a model in which suppression of ChREBP activates amino acid catabolism at a time when anaplerotic substrate influx from glucose or other sugars is limited due to impaired glycolysis. Conversely, when ChREBP is activated by carbohydrate feeding, this may serve to suppress expression of amino acid transporters to help limit gluconeogenesis from amino acids at a time when glucose is readily available. Interestingly, the transcript encoding the sodium-coupled neutral amino acid transporter-2 (SNAT2, Slc38a2), previously reported by another group to be suppressed by ChREBP^30^, was not significantly altered by GalNAc-siChREBP treatment in either the RNA-seq analysis or when measured by q-PCR in the current study. In the prior study, chromatin immunoprecipitation was used to demonstrate binding of ChREBP to the SNAT2 promoter in vivo, but suppression of SNAT2 expression by ChREBP overexpression was demonstrated only in isolated hepatocytes, and effects of ChREBP suppression were not described. It is possible that overexpression of ChREBP in liver may have resulted in similar suppression of SNAT2 expression, but this was not reported.

Knockdown of ChREBP expression in liver of HF/HS-fed rats had complex effects on lipid metabolism. Some, but not all of these effects were previously reported in studies involving partial ChREBP knockdown in a mouse model of glycogen storage disease type 1 or ASO-mediated knockdown of ChREBP in fructose-fed rats, or in other models ^4,10,17,18^. In the current study, while knockdown of ChREBP in the liver decreased DNL, it also increased total hepatic fatty acid and TG content. In addition, while increasing lipid storage in liver, knockdown of ChREBP affected indices suggestive of increased FAO, including an increase in transcripts encoding the four core enzymes of mitochondrial FAO, reduced circulating NEFA, and increased circulating ketone levels. ChREBP suppression also caused increases in a wide array of fatty acid-derived acylcarnitine species in liver. It should be noted that a study of mice with global ChREBP knockout reported reduced rates of fatty acid oxidation in perfused livers relative to wild-type control mice, measured indirectly by dilution of ^13^C-glutamate labeling^13^. A significant decrease in ketone production was also reported in that study, unlike what we observed here. Absent direct measurements of fatty acid oxidation in the current study, our best interpretation of our findings is that hepatic ChREBP knockdown results in an activation of fatty acid oxidation to the level of acetyl CoA, some of which is diverted to ketones, but the bulk of which fails to be fully oxidized due to limitations in supply of free CoA, NAD(P), and anaplerotic substrates. When viewed in this light, the accumulation of a broad spectrum of liver acylcarnitines may reflect a bottleneck in complete oxidation of fatty acids, as has also been described in skeletal muscle in obesity^26^. The apparent discrepancies with the prior study^13^ may have been related to their use of global ChREBP knockout mice in which metabolic effects of ChREBP loss would be manifest in extrahepatic tissues, versus our strategy of targeted suppression of ChREBP in liver of rats fed on HF/HS diet.

Overall, our lipid metabolism-related data provides new support for the concept that ChREBP is a transcription factor that seeks metabolic equilibrium when the system is challenged by carbohydrate feeding. When viewed in this way, increases in hepatic fatty acid and TG content observed here in response to ChREBP suppression may be counterbalanced by a decrease in DNL and an increase in fatty acid oxidation. The observed increases in hepatic TG and fatty acids under these conditions may be driven by the decrease in TM6SF2 expression and increased CD36 expression, allowing the liver to preserve lipid stores during a period of reduced fatty acid synthesis and increased fatty acid oxidation. Our comprehensive transcriptomic analysis suggests involvement of other regulatory genes in mediating this complex physiologic response to ChREBP suppression, including the decrease in expression of Rgs16, which functions as a suppressor of fatty acid oxidation, and the decrease in expression of key genes in the DNL pathway, PKLR, ACLY, and FASN. The converse of our findings on the effects of ChREBP suppression on lipid metabolism is that its activation in response to carbohydrate feeding would result in an increase in DNL, a decrease in fatty acid oxidation, an increase in lipid storage, but modulated by increased lipid export from the liver as VLDL and a decrease in fatty acid uptake.

Our interest in further investigation of metabolic regulatory effects of ChREBP in liver was stimulated in part by our recent finding that the BCKDH kinase (BCKDK) and phosphatase (PPM1K) are regulated by ChREBP, and that ACLY is an alternative BCKDK and PPM1K substrate^12^. Thus, fasting and re-feeding of Wistar rats with a high-fructose diet induces expression of ChREBP in concert with increased expression of BCKDK and reduced expression of PPM1K, and adenovirus-mediated ChREBP-beta expression in liver recapitulates the effects of fructose feeding on BCKDK and PPM1K expression^12^. In addition, overexpression of BCKDK in liver of lean Wistar rats is sufficient to increase phosphorylation of ACLY and activates *de novo* lipogenesis. Together, these observations supported an overarching model wherein feeding of diets high in fructose activate a ChREBP/BCKDK/PPM1K/ACLY/BCKDH regulatory node that contributes to increased BCAA levels and dysregulation of lipid metabolism. This mechanism may contribute to the association of BCAA with T2D^35^, as well as an association of branched chain ketoacids (BCKA) and BCKDK with NAFLD/NASH that we have recently described in human subjects^36^.

In the current study, treatment of rats with GalNAc-siChREBP caused a significant decrease in BCKDK mRNA and protein levels, consistent with our prior report of an increase in expression of BCKDK in liver in response to adenovirus-mediated ChREBP-β overexpression^12^. Here we also found that ChREBP knockdown also caused an increase in expression of the E2 component (lipoamide acyltransferase, Dbt) of the BCKDH complex as well as isovaleryl CoA dehydrogenase (Ivd), the third enzyme in catabolism of leucine, providing further evidence for a role of ChREBP in regulation of BCAA and amino acid metabolism. However, whereas GalNAc-siChREBP treatment resulted in a 75% decrease in BCKDK protein levels, this had no effect on BCKDH phosphorylation. Moreover, while GalNAc-siChREBP treatment reduced phospho-ACLY levels, this effect could be explained by a decrease in total ACLY protein.

These findings suggest that additional factors may be involved in mediating interactions between ChREBP, BCKDK, ACLY and BCKDH. Consistent with this idea, GalNAc-siChREBP treatment activated consumption of BCAA with no change in BCKDH phosphorylation or BCKA levels. A main effect of ChREBP knockdown in this context may have been to increase Dbt and Ivd expression. Our current findings may have been influenced by the multiple new metabolic effects of ChREBP suppression uncovered in this study. Also, the prior studies were performed in lean male Wistar rats or in male Zucker-obese and Zucker-lean rats fed on standard chow diet, while the current study involved obesity prone (OP-CD) male Sprague Dawley rats fed for 10 weeks on a HF/HS diet prior to treatment with the GalNAc-siRNA vectors, suggesting that nutritional status or genetics could have contributed to the discordant findings. Other recent work by our group suggests that careful attention must be paid to species (e.g. mouse versus rat versus human) and sex in analysis of this regulatory node^12,37^. For example, immunoblot analysis of livers from Zucker-lean male and female rats demonstrate that BDK expression, ACLY phosphorylation, and BCKDH phosphorylation are all higher in females than in males, suggesting that the interactions of these proteins may be regulated in a sexually dimorphic manner. We have also observed a much higher level of basal BCKDH phosphorylation in rats compared to mice, independent of diet, possibly suggesting species-dependent dynamics of BCKDK and PPM1K interactions with their substrates BCKDH and ACLY. Finally, rats, non-human primates and humans all contain a ChREBP binding sequence in the promoter region of their BCKDK genes, whereas mice lack this motif^12^. Further studies focusing on the impact of sex, species, and nutritional status on the ChREBP/BCKDK/BCKDH/ACLY regulatory node are needed.

In sum, by combining metabolomic, transcriptomic, and immunoblot analyses, we have uncovered regulatory effects of ChREBP on metabolic homeostasis far exceeding it historical role in control of core glucose and lipid metabolic pathways, to now include effects on co-factors, transporters for amino acids and other small molecules, nucleotide metabolism, and control of mitochondrial substrate supply. Future work will focus on the effects of sex and nutritional status on the regulatory networks that have emerged from these studies.

## Experimental Procedures

### Animal Studies

All procedures were approved by Duke University Institutional Animal Care and Use Committee and performed according to the regulations of the committee. Breeding pairs of Obese Prone CD (OP/CD) Sprague Dawley rats were gifts from Dr. Warren Grill and Dr. Eric Gonzalez, Duke University, and a colony was established and maintained by Duke Laboratory Animal Resources (DLAR). Starting at 4 weeks of age, male OP/CD rats were single-housed with a light cycle of 7 AM on/7 PM off, and fed *ad libitum* with a high-fat/high-sucrose (HF/HS) diet (D12451i, Research Diets) containing 47% fat (kcal) and 17% sucrose (kcal). Body weight and food intake were monitored weekly. After 9 weeks of feeding of the HF/HS diet, plasma samples were collected via saphenous vein bleeding. One week later, animals received an initial subcutaneous injection of one of two GalNAc-siRNA constructs at a dose of 9 mg/kg body weight, or an equal volume of the diluent (PBS), (see below for description of the two GalNAc-siRNA reagents). Additional doses of each GalNAc-siRNA construct were injected at 10, 18 and 25 days after the first injection. Animals were fasted overnight one day after the third injection (day 19), and subjected to an intraperitoneal glucose tolerance test (IPGTT) on the following day. Animals were weighed and a glucose solution (1g/kg body weight) was administered via intraperitoneal injection. Tail blood samples were obtained and glucose levels measured with a blood glucose meter (CVSHealth) immediately before and at 30,60, 90, 120, and 180 minutes after bolus injection of glucose. One day after the fourth GalNAc-siRNA or saline injection on day 25, plasma samples were collected via saphenous vein bleeding. A bolus of deuterium oxide (D2O, 10 ml/kg body weight, Sigma Aldrich) was then given by intraperitoneal injection and followed by free access to drinking water supplemented with 4% D2O for the rest of the experimental period. Saphenous plasma samples were collected again one day after the bolus delivery of D2O (day 27). On day 28 between 8 AM-noon, animals were anesthetized with Nembutal (250 mg/kg) by intraperitoneal injection and sacrificed for collection of plasma and liver samples that were rapidly frozen in liquid nitrogen and stored at -80°C for further analyses.

### Measurement of Hepatic Triglycerides and Glycogen

Frozen liver samples were pulverized under liquid nitrogen with a mortar and pestle. Hepatic glycogen content was measured as described previously^24^. Hepatic triglyceride content was measured using a Triglyceride Liquidcolor® kit (StanBio) after chloroform/methanol extraction^24^.

### Construction of GalNAc-siRNAs

GalNAc-siRNA reagents were prepared by Biosynthesis Inc. Each of the siRNA constructs was fully synthesized with standard 2’ modified nucleotides. Primetech TEG-GalNAc was conjugated on the 3’ end of the sense strand. Individual strands were dual HPLC purified and analyzed by mass spectrometry to verify purity of > 98% before annealing the sense and antisense strands as duplexes. The specific siRNA sequences used were: 5’CCGACCUUUAUUUGAGUCCU3’, specific for ChREBP (si-ChREBP) and 5’UUCGUACGCGGAAUACUUUCGA3’, as a non-targeting control siRNA sequence (si-Ctrl).

### Metabolomic Analyses

Amino acids and acylcarnitines were analyzed by flow injection electrospray ionization tandem mass spectrometry and quantified by isotope or pseudo-isotope dilution using methods described previously^12,28^. Levels of free coenzyme A (CoA) and short chain acyl CoAs were assayed by a previously described targeted tandem mass spectrometry method^27,38^. A panel of nucleotides, nicotinamide dinucleotide and purine pathway intermediates were measured by a previously described targeted LS/MS method^31^. Chromatographic separations and mass spec analysis were performed using a Xevo TQ-XS quadrupole mass spectrometer coupled to Acquity UPLC system (Waters, Milford, MA) and a Chromolith FastGradient RP-18e 50-2mm column (EMD Millipore, Billerica, MA, USA), and samples were spiked with a cocktail of 9 stable isotope-labeled standards to facilitate analyte quantification.

Branched-chain keto acids (BCKA) were analyzed by LC-MS/MS as previously described, using isotopically labeled internal standards KIC-d3, KIV-5C13 (Cambridge Isotope Laboratories), and KMV-d8 (Toronto Research Chemicals)^27^. Creatine, phosphocreatine, creatinine, and guanidinoacetate were analyzed by LC-MS/MS as described previously^39^. Metabolite concentrations were computed using an external calibration constructed from a serial dilution of authentic standards in dialyzed fetal bovine serum (Sigma, MO, USA). Finally, a panel of TCA cycle intermediates/organic acids were measured by targeted GC/MS analysis as described^28^.

### Conventional metabolites and hormones

Insulin in rat plasma was measured with an ELISA assay from Meso Scale Discovery (MSD, Rockville, MD). Other analytes were measured on a Beckman DxC 600 analyzer using reagents from Beckman (Brea, CA) for glucose, triglycerides, and AST, and Fujifilm Wako (Osaka, Japan) for non-esterified fatty acids (NEFA), total ketones, and 3- hydroxybutyrate.

### Measurement of hepatic *de novo* lipogenesis

*De novo* lipogenesis was measured by ^2^H2O labeling of newly synthesized fatty acids in the liver using previously described methods^12,16^. To check for enrichment of D2O in plasma water, 10 µl plasma was extracted with acetone/acetonitrile solution (1/20, volume ratio) followed by chloroform treatment and used for Gas Chromatography-Mass Spectrometry (GC-MS) analysis with an Agilent 5973N-MSD instrument equipped with an Agilent 6890 GC system, and a DB-17MS capillary column (30 m x 0.25 mm x 0.25 um). The mass spectrometer was operated in the electron impact mode (EI; 70 eV). The temperature program was as follows: 60°C initial, increase by 20°C/min to 100°C, increase by 50°C/min to 220°C, and hold for 1 min. The sample was injected at a split ratio of 40:1 with a helium flow of 1 ml/min. Acetone eluted at 1.5 min. Total palmitic acid labeling in the liver was assayed by GC-MS. Briefly, 20 mg liver tissue was homogenized in 1 ml KOH/EtOH (EtOH 75%) and incubated at 85 °C for 3 hours, and 20 µl of 10 mM [D31]palmitate was added to samples as internal standard after cool down. Extracted palmitic acid was mixed with 100 µl N-tert-Butyldimethylsilyl-N-methyltrifluoroacetamide (TBDMS) at 70 °C for 20 minutes, and the TBDMS-derivatized samples were analyzed with an Agilent 5973N-MSD equipped with an Agilent 6890 GC system, and a DB-17MS capillary column (30 m x 0.25 mm x 0.25 um). The mass spectrometer was operated in the electron impact mode (EI; 70 eV). The sample was injected at a split ratio of 10:1 with a helium flow of 1 ml/min. Palmitate-TBDMS derivative eluted at 9.7 min, and the m/z at 313, 314, and 319 were extracted for M0, M1, …… and up to M16 palmitate quantification. After correction for natural isotope abundance^40^ newly synthesized palmitic acid was calculated as: % newly synthesized palmitic acid labeling = total palmitic acid labeling /(plasma ^2^H2O labeling × 22)×100.

### Immunoblotting

Frozen liver tissue was lysed in RIPA buffer (150 mM NaCl, 50 mM Tris (pH 8.0), 0.1% SDS, 1% Triton X-100, 0.5% C24H39NaO4) with complete EDTA-free protease inhibitor cocktail (Roche) and PhosSTOP phosphatase inhibitor cocktail (Roche). 20 µg of protein were resolved by SDS-PAGE and transferred onto PVDF membranes. Membranes were blocked and probed with antibodies. Primary antibodies used were: Phospho-S293-BCKDHA (Abcam ab200577), BCKDH- E1α (CST 90198), Phospho-S454/455-ACLY (CST 4331), ACLY (CST 4332), BCKDK (Abcam ab128935), Vinculin (CST 4650), ChREBP (Novus NB400-135), and TM6SF2 antibody (pAb-505E, a gift from Dr. Helen Hobbs, UT Southwestern Medical Center, Dallas). Secondary antibodies used were: IRDye 800CW Donkey Anti-Rabbit IgG and IRDye 680RD Goat Anti-Mouse IgG (Li-Cor). Immunoblots were developed using a Li-Cor Odyssey CLx and quantified using the Li-Cor software.

### RNA isolation and qPCR analysis

RNA from rat livers was isolated using TRI Reagent (MilliporeSigma, T9424). RNA was reverse transcribed using a SuperScript VILO MasterMix (Invitrogen, Thermo Fisher Scientific). Gene expression was analyzed using QuantStudio 6 Flex Real-Time PCR System (Applied Biosystems, Thermo Fisher Scientific) and SYBR Green chemistry (PowerUp SYBR Green Master Mix; Applied Biosystems, Thermo Fisher Scientific). Gene-specific primers were synthesized by Thermo Fisher Scientific or MilliporeSigma, and a list of primers used is provided in **Table 1**. Each sample was run in duplicate. Beta-2-Microglobulin (B2m) or TATA-box binding (Tbp) transcripts was used as house-keeping genes to normalize expression levels of genes of interest.

**Table 1.**
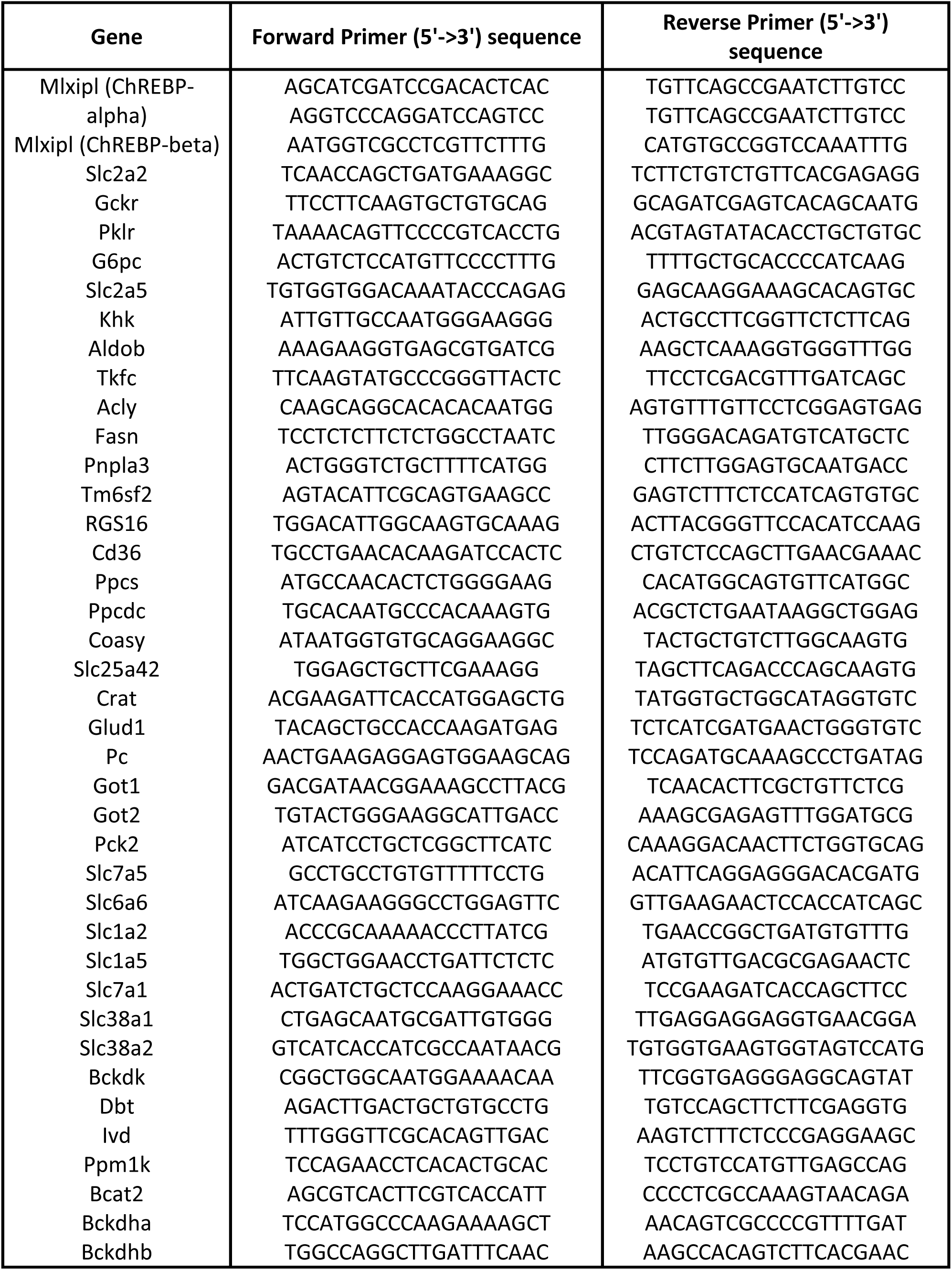
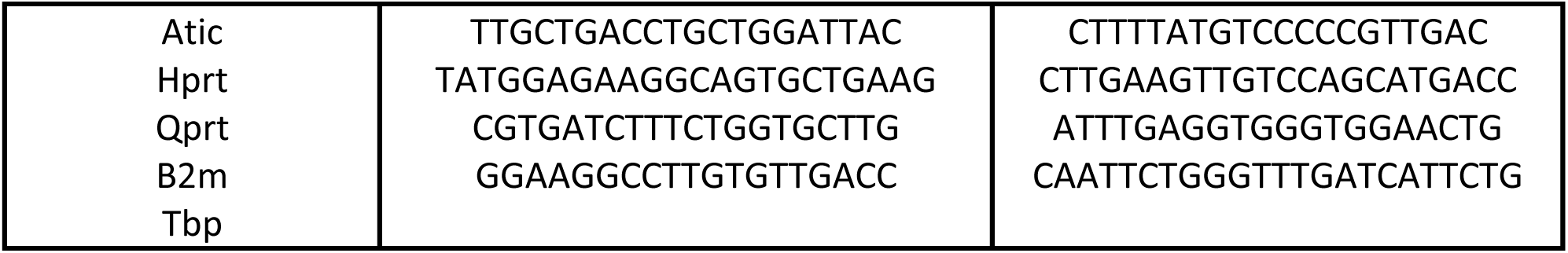
Sequences of primers used for q-PCR analyses.

### RNA sequencing and analysis

RNA was isolated from rat liver with TRI Reagent (MilliporeSigma, T9424). RNA-seq was performed by Novogene, CA. Briefly, mRNA was purified from total RNA using poly-T oligo-attached magnetic beads. After fragmentation, the first strand cDNA was synthesized using random hexamer primers, followed by the second strand cDNA synthesis using dUTP. A directional library was constructed, which included end repair, A-tailing, adapter ligation, size selection, USER enzyme digestion, amplification, and purification. 150 bp paired-end sequencing was performed on an Illumina NovaSeq 6000, and at least 77 million clean reads were obtained per sample. Pseudoalignment and quantification of transcripts was performed using Kallisto with a transcript index built from rat genome assembly mRatBN7.2^41^. Differential gene expression analysis was performed using DESeq2^42^. Gene set enrichment analysis was performed using EnrichR^43^.

### Statistical Analyses

Methods applied for statistical analysis of data are specified in Figure Legends.

## Acknowledgements

The authors are grateful to Ms. Helena Winfield for expert technical assistance in the animal studies.

## Author contributions

J. A., I.A., M.A.H., C.B.N. conceptualization; J.A., I.A., J.B., M.A.H., C.B.N. methodology; J.A., I.A., M.A.H., C.B.N. validation; J.A., I.A., M.A.H., C.B.N. formal analysis; J.A., I.A., G.Z., A.C., O.I., H.M., M.J.M., T.G. investigation; C.B.N. writing-original draft; J.A., I.A., G.Z., M.A.H., C.B.N. writing-review & editing; J.A., I.A., A.C., M.A.H. visualization; M.A.H., C.B.N. supervision; M.A.H., C.B.N. project administration; M.A.H., C.B.N., funding acquisition.

## Funding and additional information

This work was supported by National Institutes of Health grants DK12170 (to C.B.N. and M.A.H.), DK124723 (to C.B.N.), and DK100425 (to M.A.H.). We thank Dr. Anish Konkar and Eli Lilly for their gift of GalNAc-siRNA reagents used in this study.

## Supplemental Figure Legends

**Supplemental Figure 1.**
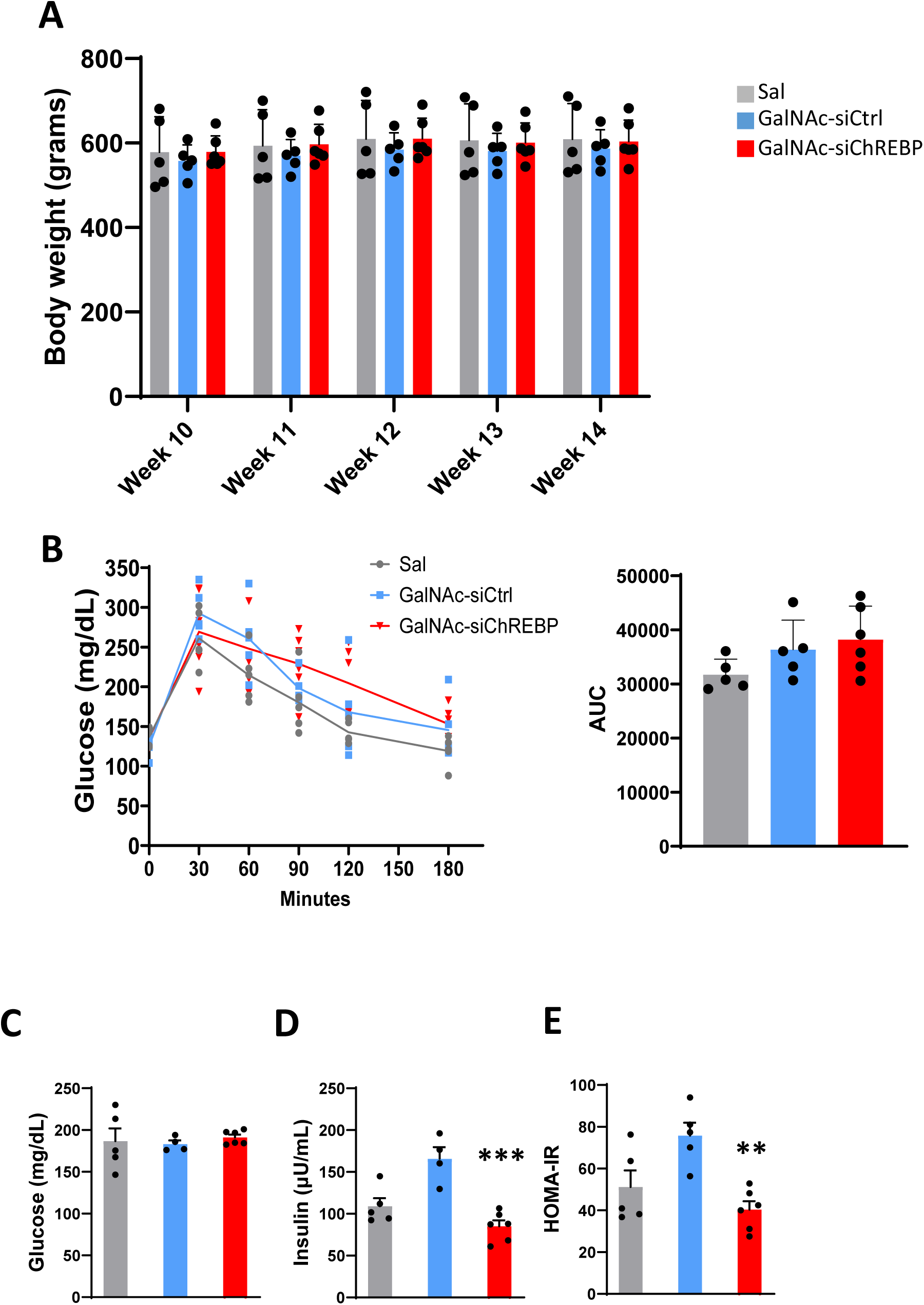
E**f**fects **of GalNac-siChREBP on body weight, interim glucose tolerance, and blood glucose and insulin levels at sacrifice.** Rats fed on HF/HS diet received multiple injections of GalNac-siChREBP, GalNac-siCtrl, or saline over a 28 day period. On day 19 after the initial injections, animals were fasted overnight and subjected to an intraperitoneal glucose tolerance test (IPGTT) via IP injection of a glucose solution (1g/kg body weight), followed by tail vein blood sampling at the indicated time points. **A.** Body weight (g) at multiple study time points. **B.** Glucose excursions during IPGTT (left) and calculated area under the curve for each group (right). **C.** Blood glucose levels at time of sacrifice (day 28 post start of GalNac-siRNA treatments). **D.** Blood insulin levels at time of sacrifice. **E.** Calculated HOMA-IR at time of sacrifice. Data are the mean ± standard error of the mean (SEM) for 6 independent rats/liver samples for the GalNac-siChREBP-treated group and 5 rats/liver samples for the GalNac-siCtrl and saline-treated groups. Statistical analysis performed using one-way ANOVA followed by Tuckey test. Symbols **, and *** indicates a measure significantly different in the GalNAc-siChREBP treatment group compared to the GalNAc-siCtrl group, with p <0.01 and p<0.001, respectively.

**Supplemental Figure 2.**
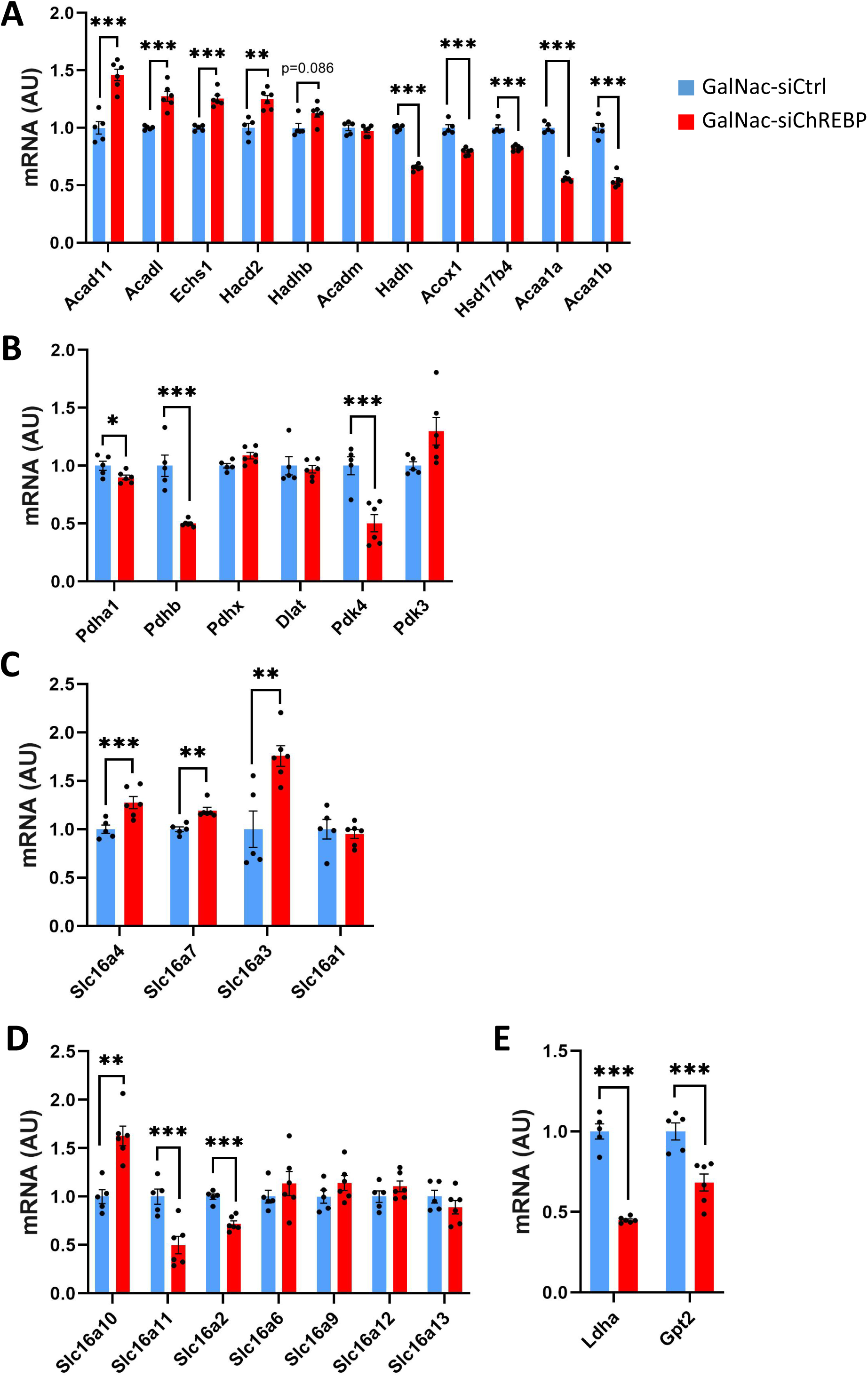
E**f**fects **of GalNac-siChREBP on selected transcript levels in liver.** Rats fed on HF/HS diet received multiple injections of GalNac-siChREBP or GalNac-siCtrl for 28 days prior to sacrifice and collection of liver samples for transcript analyses by RNA-seq. **A.** Transcripts involved in mitochondrial or peroxisomal fatty acid oxidation. **B.** Transcripts encoding components of the pyruvate dehydrogenase (Pdh) enzyme complex and its regulatory kinases. **C.** Transcripts encoding various isoforms of monocarboxylic pyruvate/lactate acid carriers. **D.** Transcripts encoding other transporters of the Slc16 family. **E.** Transcripts encoding lactate dehydrogenase a (Ldha) or alanine transaminase (glutamate-pyruvate transaminase 2, Gpt2). Data are shown as mean ± standard error of the mean (SEM). Differential gene expression analysis was performed using DeSeq2 with correction for multiple comparisons. Symbols *, **, *** indicate significant difference between groups with adjusted p-value padj< 0.05, padj < 0.01 and padj < 0.001, respectively.

